# Mapping Tumor Microenvironment and Treatment Response of Diffuse Midline Glioma Using Multiplexed Immunofluorescence and AI Models

**DOI:** 10.1101/2025.03.23.644698

**Authors:** Sandra Laternser, Andrea Joseph De Micheli, Sarah Brüningk, Elizabeth McDonough, Antonela Petrovic, Elisabeth Jane Rushing, Christine Surrette, Julia Bugajska, Augustine Eze, Lindsay Kilburn, Luisa Machado, Susanne Dettwiler, Fabiola Prutek, Noelia Casares Lagar, Daniel de la Nava, Denise Morinigo, Fiona Ginty, Matthew D. Dun, Michael Grotzer, Ana Sofia Guerreiro Stücklin, Sabine Mueller, Roger Packer, Miriam Bornhorst, Marta María Alonso, Javad Nazarian

## Abstract

**Background:** Despite its clinical promise in non-solid tumor, immunotherapy is yet to show significant clinical efficacy for brain tumors including pediatric diffuse midline glioma (DMG). This indicated the need to fully explore DMG immune tumor microenvironment (TME).

**Method:** Whole brains (49 DMGs, 20 non-DMG, 10 non-malignant) from 79 pediatric patients were used to establish a tissue microarray (918 cores) representing primary, metastatic, and adjacent healthy sites. CellDIVE MxIF multiplex assay was used to probe for 33 immune and cell type markers. RNA sequencing (n=62 patients) defined additional immune signatures. Findings were validated using patient plasma and DMG PDX models. Our annotated single-cell atlas was used to train a spatial AI model to predict antigens from H&E staining.

**Findings:** We found enrichment of M1-activated microglia in primary versus adjacent healthy tissue. PD1 positive cells were significantly (p<0.01) higher in tumor compared to adjacent controls. This was validated by mRNA profiling, further indicating two distinct groups with top 35 significant (p<0.05) genes revealing synaptic signature in the metastatic cohort.

We stratified the patient cohort by treatment. Imipridone cohort (n=5) showed decreased progenitor (Nestin+, Vimentin+, and SOX2+) and increased macrophages/microglia infiltration. Increased T and B cells was validated in patient plasma following imipridone therapy. Combination therapy of imipridone and immunotherapy (n=7) resulted in increased myeloid (Iba1, CD68, CD163) and lymphoid (CD3, CD8) cells. Enhanced immune engagement was validated in DMG PDX models. Machine learning resulted in a spatial AI model capable of predicting 22 antigens using H&E slides.

**Interpretations:** DMG tumors maintain a cold immune microenvironment, which is nevertheless dynamic and responsive to therapy, indicating the need to explore combination therapies. AI-assisted antigen detection is suitable for rapid interpretation of clinical biospecimens.

**Funding:** This work was supported by Rising Tide, SNF, LilaBean Foundation, Swifty Foundation, Swiss to Cure DIPG and Yuvaan Tiwari Foundation.

Graphical Abstract

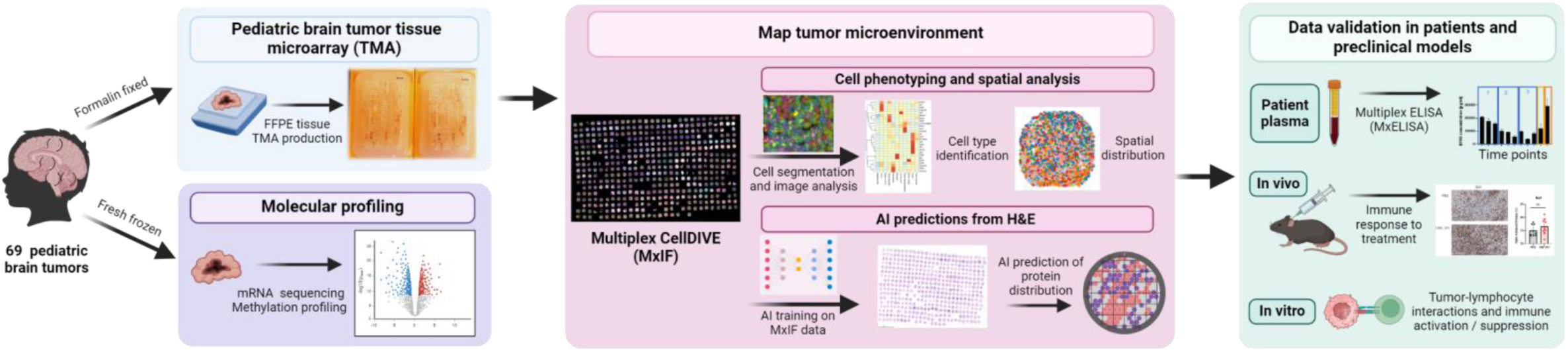

## Introduction

While advancements in pediatric oncology have improved outcomes across many childhood cancers, malignant tumors of the central nervous system (CNS) remain the main cause of cancer-related mortality in children [1]. Children diagnosed with diffuse midline glioma (DMG), including diffuse intrinsic pontine glioma (DIPG), face a particularly poor prognosis, with median survival spanning only a few months from diagnosis [2]. The challenging anatomical location and the diffuse nature of DMGs limits surgical interventions. Additionally, the ‘cold’ tumor immune microenvironment (TME) presents a significant obstacle to immunotherapeutic interventions [3]. DMGs harbor mutations that affect genes encoding H3.3 (H3F3A, 70%) or H3.1 (HIST1H3B/C, 20%) histone proteins, leading to global hypomethylation [4–6]. Despite the growing molecular understanding of these tumors, no curative treatments currently exist for patients diagnosed with DMG. Palliative radiotherapy remains the only standard of care, offering limited improvement in survival and quality of life [7, 8]. DMG TME remains poorly understood, largely due to the rarity of the disease and the limited availability of tissue biopsies and post-mortem specimens. Three studies have provided an in-depth characterization of DMG tumor microenvironment [3, 9, 10], particularly the immune cell population and their interactions with surrounding parenchyma is discussed on recent reviews [11]. This gap in knowledge extends to immune profiles across different brain anatomical regions, such as brainstem, cerebellum, thalamus and cortex.

Recent research targeting tumor metabolism [12, 13] and immunotherapeutic approaches, such as oncolytic viruses [14] have shown promising clinical efficacy, particularly by targeting proteins including GD2 or B7H3. These immune suppressive checkpoint markers are highly expressed in several tumor types [15], including DMGs [16]. Additional tumor associated immune suppressor markers including PD1 and PDL1 are scarcely expressed in DMGs as compared to other tumors [15]. To further characterize DMG microenvironment we used the multiplexed immunofluorescence (MxIF) imaging [17–20], and mapped 33 antigens representing immune and brain enriched cell types across a newly constructed TMA consisting of 816 tissue cores. The MxIF platform uses continuous staining-imaging-signal inactivation cycles to probe for unlimited numbers of antigens. This platform in combination with our TMA provided the opportunity to multiplex a large number of antigens across DMG and few other CNS tumor types.

We present multiplex imaging analyses representing immune and morphological characteristics of various brain anatomical regions across DMG and other pediatric CNS cancers. A total of 44 antibodies were used, of which 33 showed specificity. Here, we describe expression and distribution of 22 antigens across 918 tissue cores (242 FFPE blocks). We correlated immune markers with patient plasma and validated immune engagement in DMG preclinical models. Recent advances in digital pathology leveraging artificial intelligence (AI) offer a new solution to infer molecular characteristics directly from H&E sections [21]. Thus, building on digital pathology foundation models, we explore the potential of AI predictions of antigen signatures in pediatric brain tumors, demonstrating the utility of these methods in accelerating the in-depth characterization of minimally processed tissue samples.

## Results

### Establishing a pediatric brain tumor tissue microarray

We collected whole brains from 79 patients at postmortem (**Fig. 1A**), including those diagnosed with DMG (n=49), glioblastoma (GBM, n=9), ependymoma (n=5), atypical teratoid/rhabdoid tumor (ATRT, n=2), medulloblastoma (n=1), low-grade glioma (LGG, n=1), embryonal tumor with multilayered rosettes (ETMR, n=1), adult GBM (n=1), as well as non-malignant controls (n=10). We selected up to four distinct anatomical regions of the brain, including primary tumor sites, metastatic regions, and adjacent healthy tissue (**Fig. 1B-1**). Formalin fixed paraffin embedded (FFPE) blocks (224 specimens) were processed for hematoxylin and eosin (H&E) staining and reviewed by a neuropathologist to demark tumor and healthy regions. We then obtained three punch cores from each demarcated region, resulting in 918 cores, which were then aligned to generate two TMA blocks (**Fig. 1B-2**).

**Figure 1.**
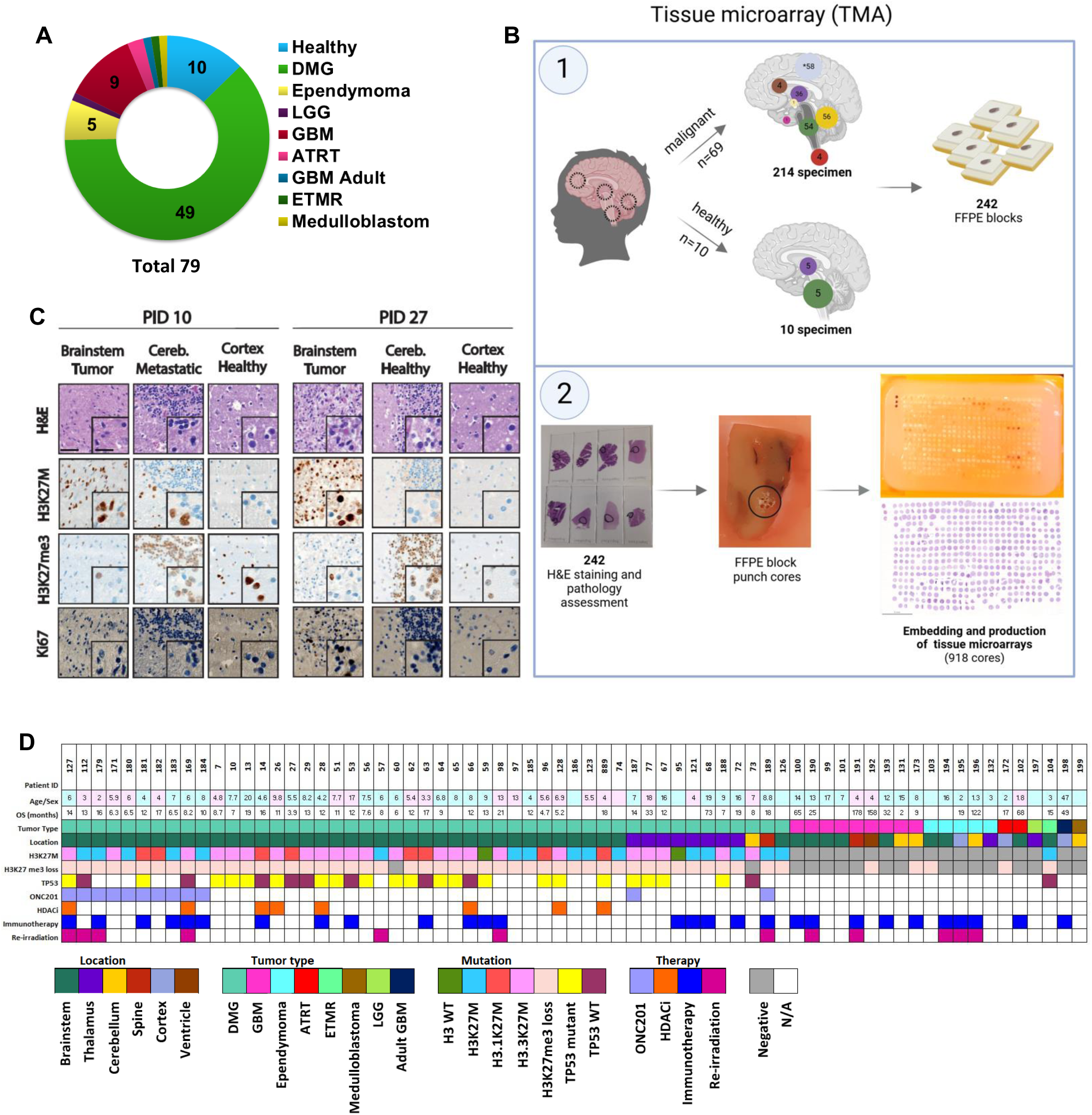
Patient cohort outline and tissue microarray production. (A) Brains from 79 patients were selected that included 49 Diffuse Midline Glioma (DMG), 9 Glioblastoma (GBM), 5 Ependymoma, 1 Low-Grade Glioma (LGG), 1 adult GBM, 2 Atypical Teratoid/Rhabdoid Tumor (ATRT), 1 Medulloblastoma, 1 Embryonal Tumor with Multilayered Rosettes (ETMR), and 10 healthy control cases. (B) A total of 224 specimens from 242 FFPE blocks representing various neuroanatomical locations from 69 brain tumor patients and 10 non-cancer controls were collected. Hematoxylin and Eosin (H&E) staining was performed to assess tumor and healthy regions by a neuropathologist. Core punches (n = 918) were obtained from pathology-designated regions and assembled into two TMA blocks (Created with BioRender.com). (C) Representative immunohistochemistry (IHC) staining of various anatomical locations representing primary, metastatic and adjacent healthy sites (two DMG patients, PID10 and PID27). Sections were probed for histone mutation maker (H3K27M), loss of trimethylation (H3K27me3), and proliferation marker (Ki67). Scale bar = 50 µm, Inset: 20 µm. (D) Oncoplot showing patient clinical, demographic, overall survival (OS), key genomic alterations and treatment information of the patient cohort used for this study.

**Figure 2.**
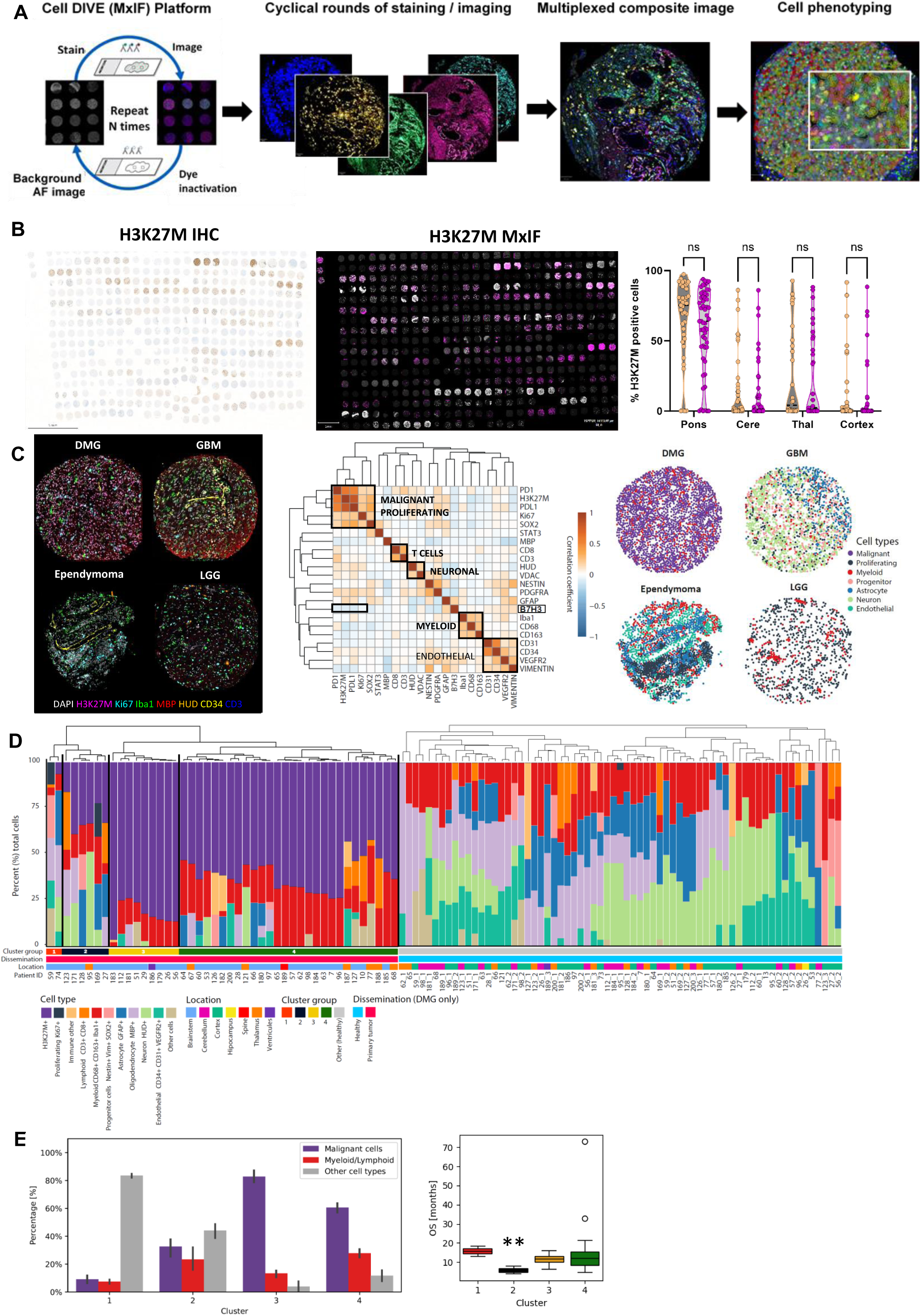
Multiplex analyses of immune marker and cell type identification. (A) The Cell DIVE (MxIF) platform employs a cyclic process of staining, imaging, and dye inactivation resulting in multiplexed immunofluorescent (IF) images, which were used for cell segmentation and phenotyping. (B) To validate MxIF, we compared IHC of H3K27M staining (left panel) with MxIF (center panel) image. Signal quantification (right graph) demonstrates the percentage of H3K27M-positive cells across four brain regions [(pons = 44), cerebellum (n = 39), thalamus (n = 29), and cortex (n = 42)]. No significant difference was observed between the two platforms (p > 0.05, 2way ANOVA, Šídák’s multiple comparisons test). (C) We probed for several antigens, representing various cell types, including DMG tumor (H3K27M), proliferation (Ki67), myeloid (Iba1), oligodendrocyte (MBP), neuronal (HUD), endothelial (CD34) and lymphocytes (CD3) using MxIF multiplexing (left panel). Hierarchical clustering (center panel) of cell type marker expression (Pearson Correlation) identified malignant, proliferating cells, T-cells, neuronal cells, myeloid cells, and endothelial cells highlighting tumor heterogeneity. B7H3 marker did not cluster with malignant cells (p < 0.0001, correlation coefficient: −0.09). Single-cell spatial distribution plots of four different brain cancer types (right panel), with each dot representing an individual cell. (D) Percentage of cell type population is provided for DMG primary sites (left) and DMG adjacent healthy (right). The four clusters identified in DMG primary sites were further interrogated to depict cell type contributions. (E) Cell type contributions (malignant, immune, other healthy tissue cells) by hierarchical clustering (left plot). OS of DMG patients stratified by the TME associated cluster type. For cluster two we observed a significantly (p < 0.01, Mann-Whitney statistical test) shorter overall survival compared to patients in other clusters (right plot). Bar plot showing mean and standard deviation (SD). (* p < 0.05, ** p < 0.01, *** p < 0.001, **** p < 0.0001)

The resulting TMA was processed for H&E and probed by immunohistochemistry (IHC) for three antigens: mutant H3K27M histone, histone 3 trimethyl mark (H3K27me3), and the proliferation marker Ki67 (**Fig. 1C, Sup. Fig. 1A**). H3K27M positivity and loss of H3K27me3 was used to score for tumor content across 558 DMG cores representing primary, metastatic and adjacent healthy sites (**Sup. Fig. 1A**). Scoring was done by a neuropathologist based on the percent of tumor cells: (0), no positive tumor cells; (1+), focal positive tumor cells; (2+), positive cells >50%. We detected H3K27M positivity in the brainstem (95%), cerebellum (51%), thalamus (48%), and cortex (frontal, occipital, temporal, and parietal regions; 14%) (**Sup. Fig. 1B**). Notably, two DMG cases were identified as H3 wild type. Clinical data including mutation burden and patient demographics are presented in **Fig. 1D**, and specimen distribution are provided in **Sup. Table 1**.

### Cell atlas of brain microenvironment using a multiplexed immuno-fluorescence platform

To establish a cell atlas, we chose 44 antigens representing immune cells, DMG biology, metabolic, and brain morphological markers **(Table 1, Sup. Table 2)**. TMA was processed for multiplex IF using the Cell DIVE™ MxIF platform which uses repeated cycles of staining and imaging and signal inactivation (**Fig. 2A**). Slides were then imaged at 20X, multiplexed composite images were captured, and cell segmentation boundaries were used to evaluate maker expression and to define cell phenotypes (**Fig. 2A**). Of the 44 antigens, 33 showed specificity, and of those, 22 were selected for further characterization for this report. To validate the MxIF platform, we first correlated H3K27M expression between our IHC staining (shown in **Fig. 1C**) and the MxIF platform. Same brain anatomical regions (pons, cerebellum, thalamus and cortex) were selected, H3K27M+ cells were counted and normalized to the total number of cells per core. Comparative analysis between IHC and MxIF showed no significant difference between the two platforms (**Fig. 2B**). We thus proceeded with generating multiplex images for analysis using the QuPath software (**Fig. 2C)** and calculated the Pearson correlation of expression patterns of co-expression between markers across all tumor types (**Fig. 2C, center panel**). We observed a similar pattern of expression amongst the following groups: malignant proliferating cells (PD1, H3K27M, PDL1, Ki67); T-cell (CD8, CD3); neuronal (HUD, VDAC); myeloid (Iba1, CD68, CD163); and endothelial (CD31, CD34, VEGFR2, VIM) **(Fig 2C, center panel)**. Surprisingly, a weak negative correlation was found between H3K27M and B7H3 **(Fig. 2C, center panel)**. The distribution of cell types across all tumors and created a spatial map highlighting the heterogeneity between and within tumor types is visualized in **Fig. 2C** right panel.

**Table 1:** Antibody panel of 44 markers used in this study.

| Classification | Antibody target | Classification | Antibody target |
| --- | --- | --- | --- |
| DMG Tumor Cells | H3K27M+, H3K27me3 loss, | Astrocytes | GFAP |
| T-Lymphocyte | CD45, CD44, CD44v6, CD4, CD3, CD8, FOXP3 | Neurons | HUD, Vimentin, NeuN, FOXG1 (neuronal development) |
| B-Lymphocyte | CD20 | Oligodendrocytes | MBP |
| Microglia / Macrophages | CD68, CD163, CD86, Iba1 | Stem cells | Nestin, SOX2 |
| Immune evasive cells | CD47, PD1, PDL1, B7H3 | Proliferating cells | Ki67 |
| Vascularization | VEGFR2, CD31, CD34, PDGFR-alpha | Pathway regulator | Akt, mTor, 14-3-3 sigma, pERK |
| Cell metabolism | VDAC1, FOXO1, Stat3 | Apoptosis | Beclin1, Foxo3a |
| Hypoxic cells | HIF1-alpha | Cell boundry | Beta-actin, NaK ATPase |
| Cell nucleus | DAPI |  |  |

**Table 2:** Antibody panel of 22 markers and their boundary used for cell segmentation.

| Marker | Boundary | Marker | Boundary | Marker | Boundary | Marker | Boundary | Marker | Boundary | Marker | Boundary |
| --- | --- | --- | --- | --- | --- | --- | --- | --- | --- | --- | --- |
| H3K27M | Nucleus | Iba1 | Nucleus | CD3 | Nucleus | B7H3 | Nucleus | PDL1 | Cell | VDAC | Cell |
| Ki67 | Nucleus | CD68 | Nucleus | GFAP | Cell | NESTIN | Cell | STAT3 | Cell | CD34 | Cell |
| SOX2 | Nucleus | CD163 | Nucleus | VEGFR2 | Cell | VIMENTIN | Nucleus | PD1 | Nucleus |  |  |
| CD8 | Nucleus | CD31 | Cell | HUD | Cell | PDGFRA | Cell | MBP | Cytoplasm |  |  |

To phenotype cell populations across the entire TMA, we performed unsupervised Louvain clustering after applying Harmony batch correction for each TMA core triplicate. We identified 10 distinct cell types including H3K27M+, proliferating, immune (myeloid and lymphocytes), progenitor, astrocytes, oligodendrocytes, neurons, and endothelial cells (**Fig. 2D**). Uniform Manifold Approximation and Projection (UMAP) analysis, clustered H3K27M+ and proliferating cells, separately from neurons, healthy astrocytes, and oligodendrocytes cluster **(Sup. Fig. 2A)**. We then focused on primary DMG (**Fig. 2D** left) and their adjacent healthy specimen (**Fig. 2D** right). The adjacent healthy tissue was enriched with a diverse cell population including neurons, astrocytes, progenitor cells and oligodendrocytes (**Fig. 2D**).

DMG primary tissue consisted of H3K27M+ malignant, myeloid, oligodendrocytes, and endothelial cells (**Fig. 2D**). DMG primary specimen further fell into four subgroups (clusters 1-4). We interrogated cell composition between these groups and found DMG primaries varied based on the presence of H3K27M+ malignant and other (healthy / non-immune) cell types (**Fig. 2E**, left plot). To understand the clinical relevance of this, we obtained overall survival for these patient population and found that patients in cluster 2 with a significantly (p<0.05) lower survival when compared to other clusters.

When comparing patient overall survival (OS) between clusters in DMG patients (n = 37) with available follow-up clinical information, we found a significantly (p = 0.008) lower OS of cluster two patients (**Fig. 2E, right plot**). These patients displayed an equal composition of malignant, immune, and healthy cells. No correlation between the fraction of malignant cells and OS was observed in any cluster (**Sup. Figure 2B, top**). For myeloid cell fractions, only in cluster 3 we observed a significant (p=0.002) correlation of the myeloid cell fraction and OS (**Sup. Figure 2B, bottom**). We further investigated these clusters in the context of treatment (ONC201, HDAC inhibitors), we found no clear distinction within the clusters. Comparing the clusters by location revealed no difference between brainstem (n = 36) and thalamic (n = 7) DMGs **(Sup. Fig. 2C**). Supplementary Table 1 lists the number of patients, specimen, FFPE blocks, punch cores, and markers, which were analyzed with MxIF staining **(Sup. Table 1)**.

### Antigen mapping across primary DMG and other CNS tumor types

We analyzed the immune landscape across all primary tumor types. Comparisons were done with DMGs as one group, and all other CNS primary sites (GBM, ependymoma, medulloblastoma, ATRT, ETMR, and LGG) as the second group. Adjacent healthy and non-malignant brain specimens were used as control. Immune cell profiling was based on the positivity threshold of the following antigens: T-cells (CD3+, CD8+), microglia (CD68+, Iba1+), and immune suppressive markers (CD163+, B7H3+, PD1+, PDL1+) **(Fig. 3A)**. Percent positivity of each antigen was calculated as the ratio of positive cells to the total number of cells **(Fig. 3A, IF upper panel and quantification lower panel)**. We observed a significantly (p < 0.05) higher CD3+ in control when compared to DMG. A small percentage (<5%) of all cells exhibited CD8 positivity which was comparable between tumors and control tissue (**Fig. 3A)**. Macrophages and microglia cells (CD68+ and Iba1+) consisted of a larger cell population (mean = 11.5% for CD68, mean = 16.5% for Iba1) across all specimens, while CD68 was significantly (p < 0.05) higher expressed in other tumors compared to control. The immune suppressive marker CD163 was consistently expressed across all tumor types with the highest percentage (>40%) in other tumor category. The percentage of PD1+ cells was significantly higher (p < 0.05) in DMG (mean = 2.4%) than other tumors (mean = 0.4%). PDL1+ cells were significantly (p < 0.001) higher in the DMG versus other tumors and control (mean DMG = 37.7%; other tumors = 9.1%, control = 1.9%). Notabley, three DMG patiens exhibited PDL1 positivity of over 90%. B7H3 (CD276), an immune suppressive marker, was detected in both DMG (mean = 3.6%) and other cancers (mean = 2.3%). Surprisingly, B7H3 expression was not significantly higher DMGs compared to healthy non-cancer brain specimens that showed highest percentage of positivity 24%.

**Figure 3.**
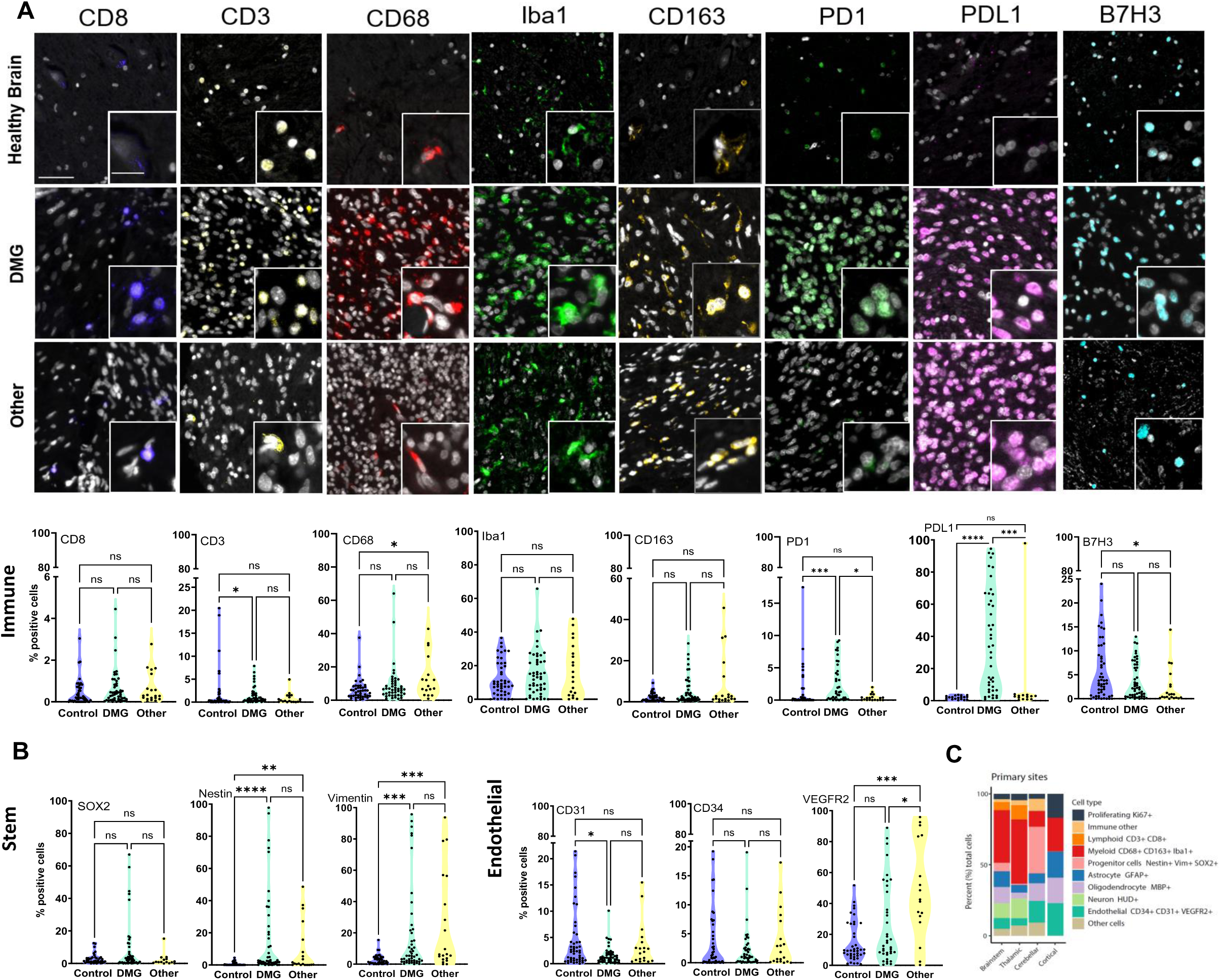
Immune, endothelial, and stem cell landscape across pediatric brain tumors. (A) Expression of eight immune markers were probed using the multiplex platform across primary sites of DMGs and other brain tumor types as well as healthy non-cancer brains. Representative IF images and quantification of number of positive cells for each specific marker across all samples are presented as violin graphs. (B) We then probed for cells expressing stem cell (SOX2, Nestin, Vimentin), and endothelial markers (CD31, CD34, VEGFR2) across the same cohort. (C) Percentage cell type composition across primary sites of all tumor types. Brainstem and thalamic primary tumors composed of the largest immune cell population. Scale bar = 50 µm, Inset: 20 µm. Kruskal-Wallis test for non-parametric data for multiple group comparisons. Dunn’s post-hoc test was used for pairwise comparisons. (* p < 0.05, ** p < 0.01, *** p < 0.001, **** p < 0.0001)

Next, we evaluated markers of stemness (SOX2, nestin, vimentin) and endothelial (CD31, CD34, VEGFR2) markers **(Fig. 3B)**. Across tumor types, DMGs displayed the highest (up to 90%) positivity for stemness antigens. Nestin and vimentin expression were significantly (p < 0.001) higher when compared to controls (**Fig 3B)**. Endothelial markers CD31, CD34 and surprisingly VEGFR2+ cells were not enriched in DMGs compared to control and other brain cancers **(Fig. 3B)**. The average VEGFR2 expression was 48.8% (others), followed by 26% (DMGs), and 15% (control).

Comparing primary sites of different tumor types, again DMG primary sites (brainstem & thalamus) comprised a large proportion of myeloid proportion compared to cerebellar and cortical tumors that were enriched in endothelial cell phenotypes **(Fig. 3C)**.

To assess immune, stem and endothelial landscape across primary tumor sites, we compared tumors from each primary site (brainstem, thalamus, cerebellum, and cortex) to neuroanatomically matched healthy tissue **(Sup. Fig. 3A)**. T-cell antigens (CD3 and CD8) were generally low (7.8% for CD3 and 4.5% for CD8) across all primary sites. Again, B7H3 positivity was equally enriched in tumor sites as compared to healthy sites. However, CD68+ (mean = 27.7%) and CD163+ (mean = 15%) cells in primary cortical tumors were significantly more (p < 0.05) compared to healthy tissue. PD1 positivity was significantly higher (p < 0.05, mean = 2.3%) in primary brainstem. PDL1+ cells were significantly (p < 0.01) higher (mean = 29.8%) in primary thalamic sites compared to healthy thalamic specimens.

Brainstem tumors were significantly (p < 0.05) enriched in SOX2 expressing cells. Vimentin was significantly higher (mean = 44.2%, p < 0.01) in primary cerebellum and primary cortical (mean = 46.3%, p < 0.05) sites. Nestin was significantly higher in primary thalamic (mean = 11.2%, p < 0.01) cerebellum (mean = 7.8%, p < 0.05), and cortical (mean = 28.8%, p < 0.01) sites. Endothelial markers (CD31, CD34 and VEGFR2) were not differentially expressed amongst this cohort, with the exception of VEGFR2 expression in primary cerebellar and cortical primary sites which were significant (mean = 70.3% and 57.9%, p < 0.05 and p < 0.01, respectively) compared to adjacent healthy sites **(Sup. Fig. 3A).**

### Antigen expression across DMG primary and metastatic sites

To assess the immune microenvironment across primary and metastatic sites, we analyzed different metastatic sites for immune cell infiltration. Specimens were reviewed and scored by a neuropathologist based on H3K27M abundance to establish the following groups: Met+3 (>50% H3K27M+ cells), Met+2 (20 – 50 %), Met+1 (5 - 20%). Metastatic sites were then processed for quantification of cytotoxic T-cells (CD8), myeloid cells (CD68), tumor-associated macrophage (CD163), and immune suppressive T-cell markers (PD1) **(Fig. 4A)**. PD1+ cells were significantly (p < 0.01) enriched in metastatic (Met+2 mean = 2.4%) and primary sites (mean = 2.6%) compared to control (mean = 0.2%) and adjacent healthy (mean = 1.3%). CD8, CD68, and CD163 were higher in tumor sites compared to control but not significantly (**Fig. 4A**). Primary sites were significantly (p < 0.05) enriched in M1 microglia (CD68+, Iba1+, CD163-, mean = 3.5%) compared to M2 (Iba1+, CD68+, CD183+, mean = 1.5%), resident microglia (Iba1+, CD68-, CD163+, mean = 1.3%), and macrophages (Iba1-, CD68+, CD163+, mean = 1.1%) (**Fig. 4B**). Analyses across primary and metastatic sites showed decreased M1 microglia abundance correlating with decreased H3K27M+ tumor cell content. M1 microglia were significantly (p < 0.0001) higher in primary compared to adjacent healthy sites (mean = 0.3%) (**Fig. 4B**). We detected only a significant (p < 0.05) expression of two stem cell markers (nestin and vimentin) in primary versus metastatic or adjacent healthy sites (**Sup. Fig. 3B**), but no significant difference in expression of endothelial (CD31, CD34, VEGFR2) markers across primary or metastatic sites (**Sup. Fig. 3B**).

**Figure 4.**
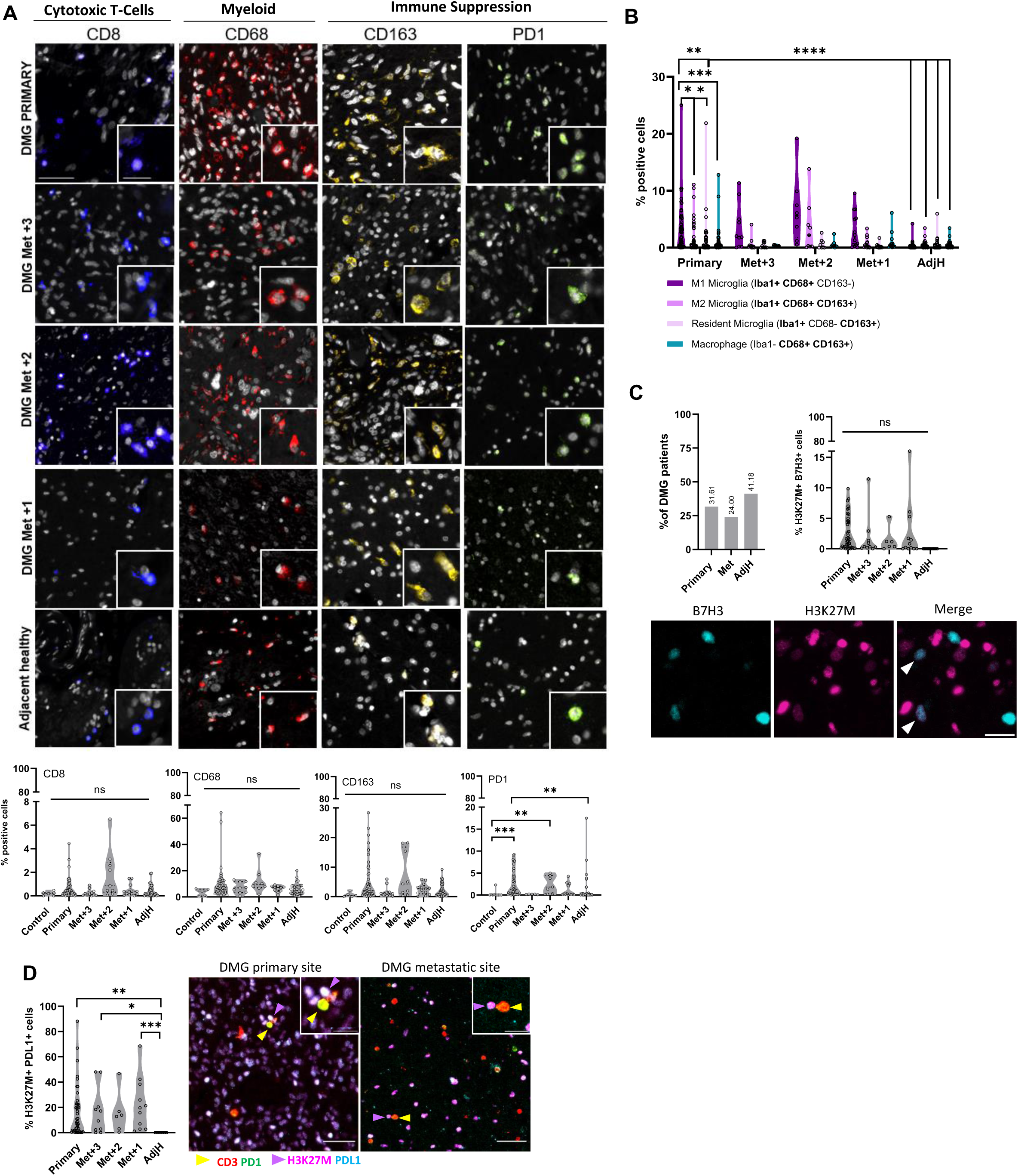
Comparative analyses of immune landscape and cellular composition across DMG primary and metastatic sites. (A) To define immune engagement in DMG, we probed for immune cell phenotype markers for cytotoxic T-(CD8), myeloid (CD68), and immune suppressive (CD163 and PD1) cells, in primary, adjacent healthy, and three metastatic sites (Met+3, Met+2, Met+1). IF (top panel) and percentage cells expressing each marker (lower panel) are shown. Myeloid and T-cells were equally distributed amongst primary and metastatic sites (p > 0.05), whereas PD1 positive cells were significantly higher in primary (p < 0.01) and Met+2 (p < 0.01) sites versus healthy control tissue. (B) Quantification of microglia and macrophage population in primary and metastatic DMG sites were done using markers of M1 activated microglia (CD68+, Iba1+, CD163-), M2 activated microglia (CD68+ Iba1+ CD163+), resident microglia (CD68-, Iba1+, CD163+), and macrophages (CD68+, Iba1-, CD163+). M1 microglia were significantly (p < 0.05) more abundant in primary sited compared to M2 microglia and when compared to adjacent healthy sites (p < 0.0001). (C) B7H3 positivity was assessed in all DMG primary, metastatic and adjacent healthy sites. A patient sample was deemed positive if B7H3 was detected in > 5% of total cells counted (left panel). B7H3 positive cases were probed for co-expression of B7H3 and H3K27M (quantified in middle panel and visualized in the right panel). (D) We counted H3K27M and PDL1 double-positive cells in primary, metastatic, and adjacent healthy tissue, to prob for T-cell interaction with tumor cells. Primary (p < 0.01) and metastatic sites showed significantly (p < 0.05 and p < 0.001) higher double positive cells when compared to adjacent healthy sites but not between each other (p > 0.05) (left panel). T-cell (PD1+ and CD3+) direct engagement with DMG cells (H3K27M+ and PDL1+) was detected in primary and metastatic brain tissue (right panel). Scale bar = 50 µm outer panel, and 20 µm in insets. Kruskal-Wallis test for non-parametric data for multiple group comparisons, and Dunn’s post-hoc test for pairwise comparisons. 2way ANOVA for parametric data and Tukey’s post-hoc test was applied for pairwise comparisons. (* p < 0.05, ** p < 0.01, *** p < 0.001, **** p < 0.0001)

### B7H3 abundance is not specific to tumor sites and localizes to H3K27 wildtype cells

Given the surprising observations about B7H3 across tumor types (**Fig. 3A**) we revisited our DMG cohort (n = 49 patients) and found B7H3+ cells only in 31% (n = 15 patients) of cases. B7H3 was detectable in only 24% (6 of 25) patients with metastatic sites (**Fig. 4C**). Adjacent healthy from 41% of patients (n = 14 of 34) showed positivity for B7H3. We then investigated the number of cells double positive for B7H3 and H3K27M. We found a modest percentage of cells to be double positive in primary (mean = 2.5%, with highest 9.8%) and metastatic sites (mean = 1.9%, with highest 16%). No double positive cells were detected in adjacent healthy sections (**Fig. 4C**). Immunofluorescent imaging confirmed this showing minimal co-expression between H3K27M+ and B7H3+ cells, whereas only weak H3K27M+ cells were B7H3+ (**Fig. 4C**, IHC). To validate this, we obtained fresh tumor and adjacent healthy specimen from a patient with DMG at autopsy. Our FACS analyses probing for B7H3+ cells showed similar B7H3+ cells in tumor (59%) and healthy (41%) specimen validating our histological analyses **(Sup Fig. 3C)**. Finally, given that B7H3 is a tumor suppressor, we investigated whether B7H3 expression in 15 patients resulted in decreased survival and did not detect a significant difference between these and B7H3 negative cohort (**Sup. Fig. 3C**).

### PDL1+ DMG tumor cells interact with PD1+ immune cells

We analyzed co-expression of PDL1 and H3K27M in DMG primary, metastatic and health sites. We found a significant (p < 0.05) number of cells co-expressing PDL1 and H3K27M in all DMG compared to healthy sites (**Fig. 4D**). Primary DMGs were enriched significantly (p < 0.01, mean = 15.7%) in double positive cells with a range up to 80% of H3K27M cells showing PDL1 expression (**Fig. 4D**). Close proximity Physical interactions between PD1+ CD3+ T-cells and H3K27M+ PDL1+ tumor cells were detectable in primary and metastatic sites (**Fig.4E, right IF images**)

### Transcriptome landscape across primary and metastatic tumors

To expand on the antigen mapping capacity of MxIF, we performed bulk RNA sequencing on frozen specimens corresponding to TMA samples. This included primary (n = 59) and metastatic tumors (n = 28), comprising of DMGs (n = 47) and other tumor types (n = 15) (**Fig. 5A**). UMAP projection and clustering of transcriptomic profiles revealed two distinct groups (A and B) (**Fig. 5B, C**). Notably, these clusters did not correlate with dissemination status (primary vs. metastatic), ONC201 or immune therapy treatment, or anatomical location—except for cerebellar samples (**Fig. 5B**). Furthermore, no association was observed between transcriptional clustering and cell type composition groups defined by our spatial analysis (**Fig. 2D**). These findings suggest that sample clustering reflects broader transcriptional programs that transcend local cell type composition, with RNA-protein expression discrepancies potentially contributing to the observed differences.

**Figure 5:**
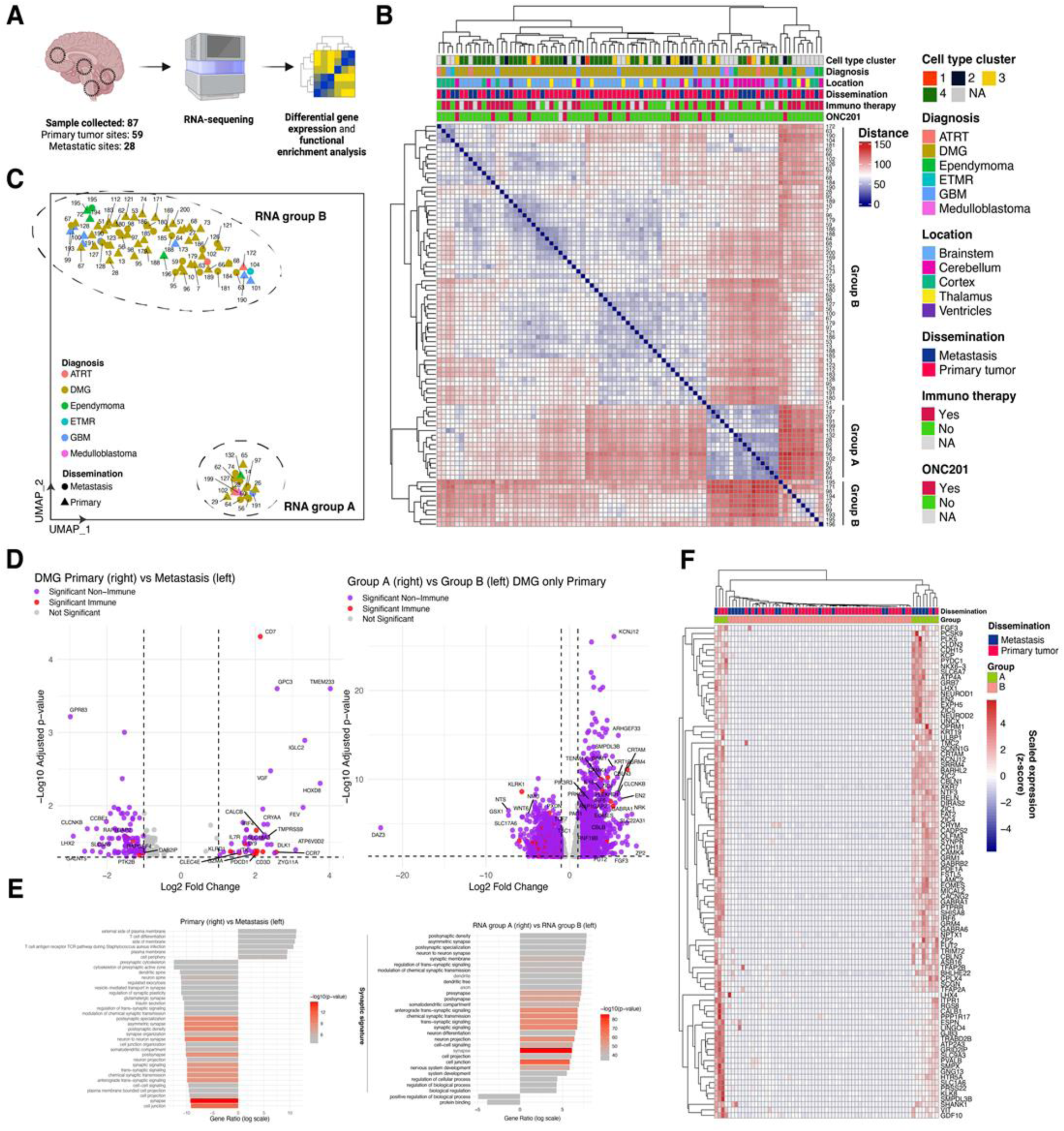
The transcriptomic landscape of DMGs and other pediatric brain tumors. (A) RNAseq workflow. Brain specimens (n = 87) were collected from 59 primary tumors and 28 metastatic sites, including samples from DMG (n = 47) and other brain tumors (n = 15). RNA sequencing was performed to analyze differential gene expression and conduct functional enrichment analysis (Created with BioRender.com). (B) Heatmap showing hierarchical clustering of samples based on normalized gene expression profiles, with annotations for cell type clusters from TMA phenotyping (see Fig. 2E), diagnosis, tumor location, dissemination status, immunotherapy, and ONC201 treatment group. Samples clustered into distinct groups (group A and B) as confirmed by UMAP projection (C).(C) UMAP plot displaying two distinct RNA groups (A and B). Samples are annotated by diagnosis type and dissemination status, showing segregation of samples into two transcriptionally distinct groups. (D) Volcano plot of differential gene expression comparing DMG primary vs metastasis sites (left panel), and RNA groups A vs B (right panel) as defined in (C). Significance (a<0.05) and fold-change in expression represented a log10 and log2 scale, respectively. Immune-related genes (cf. methods) colored in red. (E) Functional enrichment analysis (g:Profiler) from differentially expressed genes identified in (D) comparing DMG primary vs metastasis sites (left panel) and RNA group A vs B (right panel) Top 35 significant (a<0.05) gene ontology (GO) terms are shown, revealing a synaptic signature in metastatic and group A cases. Positive gene ratio terms correspond to positive log2 fold-change genes in (D) and group. Absolute higher gene ratio indicates stronger GO term association. GO terms are colored by enrichment significance on a log scale. (F) Heatmap showing scaled (z-score) expression of significantly enriched and expressed synaptic genes across dissemination (primary vs. metastasis), as well as group A and B.

We next examined differential gene expression programs across primary and metastatic tumors, treatment status, and between group A and B. Primary tumors exhibited higher expression of immune-related genes compared to metastatic sites (**Fig. 5D**, left panel), with the lymphocyte marker CD7 emerging as the most significantly upregulated gene. Functional enrichment analysis of primary tumors revealed a significant enrichment for T-cell-associated pathways, whereas metastatic tumors displayed upregulation of neuronal and synaptic mechanisms (**Fig. 5E**, left panel). When comparing RNA profiles of DMG samples from patients treated with ONC201 versus treatment-naïve individuals, we observed only minimal transcriptional differences. Notably, untreated individuals were enriched by GO terms of synaptic processes, a feature also measured in metastatic samples. Additionally, there were no differences in immune gene expression between treatment groups, suggesting that ONC201 treatment does not significantly alter the immune transcriptomic signature within DMG samples. These findings are consistent with antigen staining observations obtained via MxIF (**Sup. Fig. 4A**). Finally, we compared RNA profiles between Groups A and B. Group A samples were enriched for synaptic genes (**Fig. 5E**, right panel and **5F**), similar to untreated patients and metastatic samples. These genes include NEUROD1 and NEUROD2, transcription factors essential for neuronal differentiation and synaptic plasticity, and SYNPR, as well as genes from the GABA-A receptor family (GABRA1, GABRA6, and GABRB2) involved in inhibitory neurotransmission. These findings suggest that group A cases exhibit a transcriptional program promoting synaptic connectivity and function, distinct from other DMG subgroups. To rule out the influence of cell type composition on these synaptic signatures, we compared the cell type content of matching cores between the two groups and found no significant differences (**Sup. Fig. 5C**). This supports the hypothesis that the observed synaptic enrichment in group A, similar to that of metastatic and drug-naive patients, is driven by transcriptional programs from the cancer cells themselves rather than variations in cell type composition.

### Changes in the immune landscape in response to therapy

We further investigated the effect of treatment on the tumor microenvironment in DMG patients and stratified our patient cohort by patients who had received immunotherapy (Newcastle disease virus [22] (n = 9), treated with ONC201 (n = 5), combination of immunotherapy and ONC201 (n = 7) and ONC201 and immunotherapy naïve patients (n = 17). Progenitor, astrocytes, oligodendrocytes, neurons were enriched in immunotherapy, ONC201, treatment naïve, and combination therapy respectively (**Fig. 6A**). Myeloid (Iba1, CD68, CD163), lymphoid (CD3, CD8) cells were highly presence in combination and ONC201 treated tissue indicating treatment induced immune activation. No differential expression of other immune markers was detected (**Sup. Fig. 4A**). Combination therapy resulted in (non-significant) increase in B7H3, Iba1, and activated microglia (**Sup. Table 3**). Other clinical interventions including HDAC inhibitors (Panobinostat, Vorinostat etc.) and chemotherapy did not result in a significant change in immune composition (**Sup. Fig. 2**).

**Figure 6:**
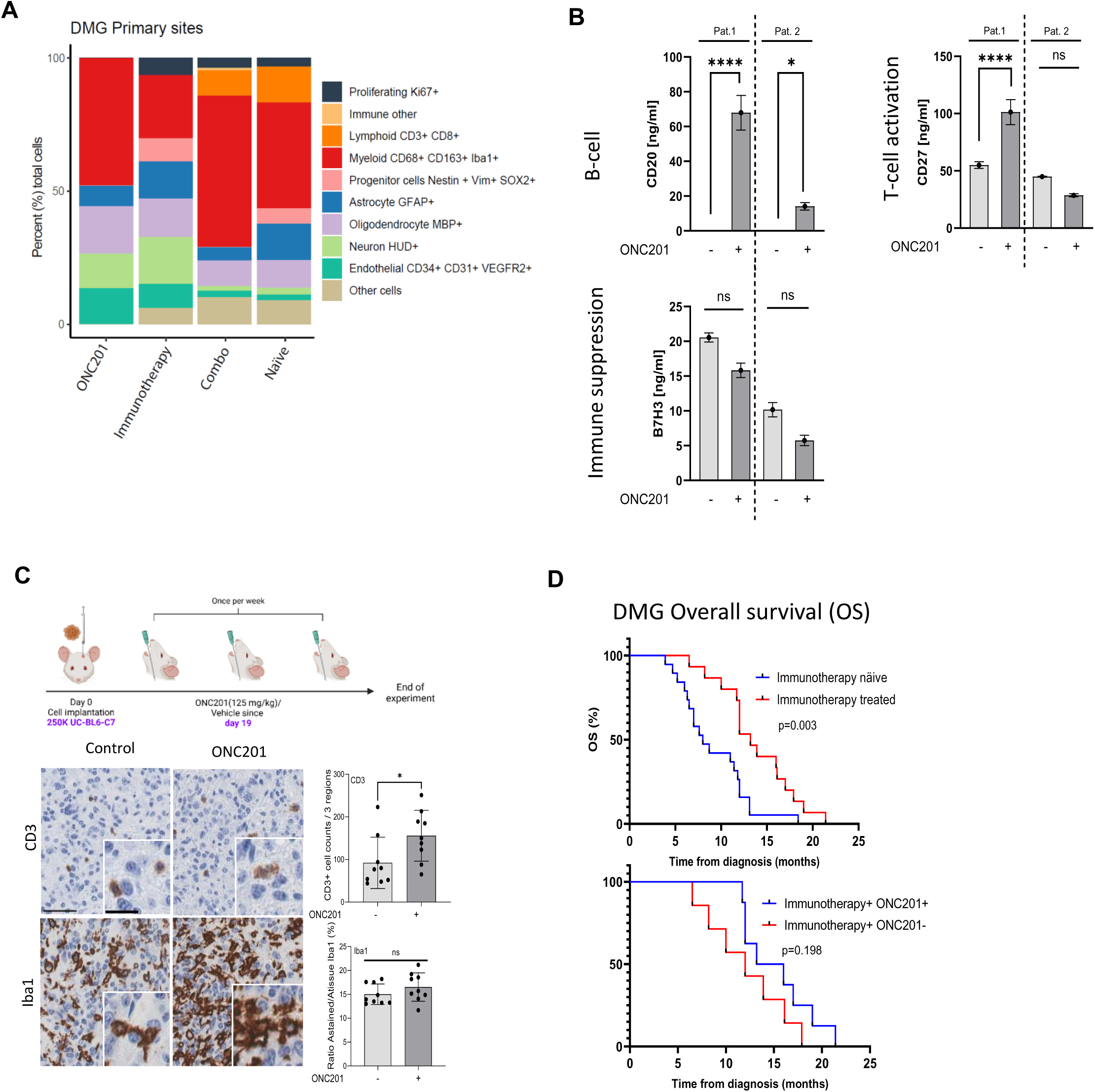
Immune engagement in the context of treatment modality. (A) Cell type composition (%) in patients treated with ONC201 (n = 5), immunotherapy (n = 9), combination therapy (n = 7), and (ONC/Immunotherapy) naïve patients (n = 17) highlight the distribution of cell populations across different treatment groups. (B) Plasma from patients receiving ONC201 were collected pre and post therapy and assessed for B-cell, T-cell, and immune suppression marker B7H3. Bar plot showing mean and standard deviation (SD). (C) DMG PDX models were treated with ONC201 (125mg/kg, 1/week) 19 days post-tumor cell implantation. Tissue was harvested and probed by IHC for expression of immune marker CD3 and Iba1. ONC201 resulted in increased CD3 expression (fold change 1.7, p < 0.05). Bar plot showing mean and standard deviation (SD). (D) Overall Survival (OS) of immunotherapy (n = 16) treated and immunotherapy-naive DMG patients (n = 20) indicated a significant increase in OS (p = 0.003, 3 months). Amongst the immunotherapy treated patient cohort, those who had received ONC201, showed an increased (3.5 months) but not significant (p = 0.2) OS. Scale bar = 50 µm outer panel, and 20 µm in insets. Statistical analysis for overall survival plots was performed using Log-rank (Mantel-Cox) test, for plasma analysis a 2way ANOVA and Tukey’s post-hoc test was applied for pairwise comparisons and for PDX IHC analysis an Unpaired t test. (* p < 0.05, ** p < 0.01, *** p < 0.001, **** p < 0.0001)

To further validate the effect of the immune signature after treatment, we used plasma from two patients receiving ONC201 and assessed the abundance of several immune markers using the sensitive MesoScale Discovery (MSD) platform. We found a non-significant decrease of immune suppression markers B7H3 after ONC201 treatment. However, we detected a significant increase in one patient for T-cell marker CD27 (FC = 1.8; p < 0.0001) and in both patients for B-cell marker CD20 (FC = 67.9 and 14.1; p < 0.0001 and p < 0.05) (**Fig. 6B**).

To validate these observations, we treated immune competent DMG murine models with ONC201 and found a significant increase (FC = 1.7; p < 0.05) in T-cell (CD3) infiltration in ONC201 treated mice (**Fig. 6C**).

To investigate the clinical impact of immune activation in patients, we collated survival data from our patient cohort and found that a significant (p < 0.003) increased (3 months) survival of patients receiving immunotherapy (**Fig. 6D**). Combination of immunotherapy and ONC201 resulted in non-significant increased survival (**Fig. 6D**).

### AI-assisted biomarker prediction from H&E

Finally, given the insight generated by spatial proteomic characterization, we provide a novel computational pathology pipeline to translate these assessments to minimally processed H&E samples. The data generated within our TMAs covers a unique variety and number of pediatric brain tumor samples, and particularly DMGs, allowing for a data-driven identification of the presence or absence of cells expressing target proteins from H&E imaging. Importantly, despite the unique size of the available data, we harness state-of-the-art foundation models to extract meaningful representations from the raw H&E images, building on heavily pretrained architectures to account for inherent challenges of data scarcity and variability across sources. We benchmarked six approaches for feature extraction including established pipelines building on Res-Nets and recent publications harnessing Vision Transformers trained on up to 1.48 million whole slide images [23–27].

We successfully trained (n = 22, one per marker) independent prediction models as a combination of these feature extractors and a light GBM classifier of specific cell types and biomarkers on the co-registered multiplex IF and (virtual) H&E imaging data (**Fig. 7A**). This approach predicts the presence of marker positive cells within image tiles of 100×100µm edge length. Importantly, we trained all models exclusively on TMA 1 data allowing us to report prediction performance on an external test set comprising 405 cores from TMA 2. We observed a performance advantage for feature extractors building on vision transformers (Virchow, Prov-GigaPath, and UNI models) with the top pipeline building on UNI, achieving competitive performance for the majority of selected markers (independent test set ROC AUC 0.57 to 0.90, **Fig. 7B**). This demonstrated the feasibility of extracting in-depth tissue characteristics and insight into the TME from H&E images in the absence of staining. In Fig. **7C** we highlight the performance based on UNI feature extraction in terms of ROCAUC and the relative area under the precision recall curve to account for the setting of imbalanced class labels. In particular, the detection of H3K27M+ DMG cells (ROCAUC_H3K27M_ = 0.88+\-0.01) as well as nestin+ neural stem/progenitor cells (ROCAUC_NESTIN_ = 0.86+/- 0.01) and MBP (MBP, ROCAUC_MBP_ = 0.90+/- 0.01) was highly robust. While immune cell markers such as CD3, CD8 or CD68 could not be detected reliably from virtual H&Es, immune suppressive signals such as PD1 and CD163 showed moderate performance (ROCAUC_PD1_ = 0.71+/-0.01, ROCAUC_CD163_ = 0.75+/-0.01). In **Fig. 7D** we provide spatial prediction examples in selected heterogenous (and hence particularly challenging) tissue cores for H3K27M, PDGFRA, PD1, and CD34. We further analyzed the prediction performance in sample subsets of the independent test set to validate the generalizability of our models to different tumor types. **Figure 7E** shows the independent test set ROCAUC for all other primary tumors. While we obtained similar performance for most of the markers for DMG and other primary tumor samples, H3K27M was poorly predicted for other type of tumors (note the absence of any positive instances here). As such our data support the utility of AI for predicting tumor microenvironment which may be complementary to current platforms.

**Figure 7:**
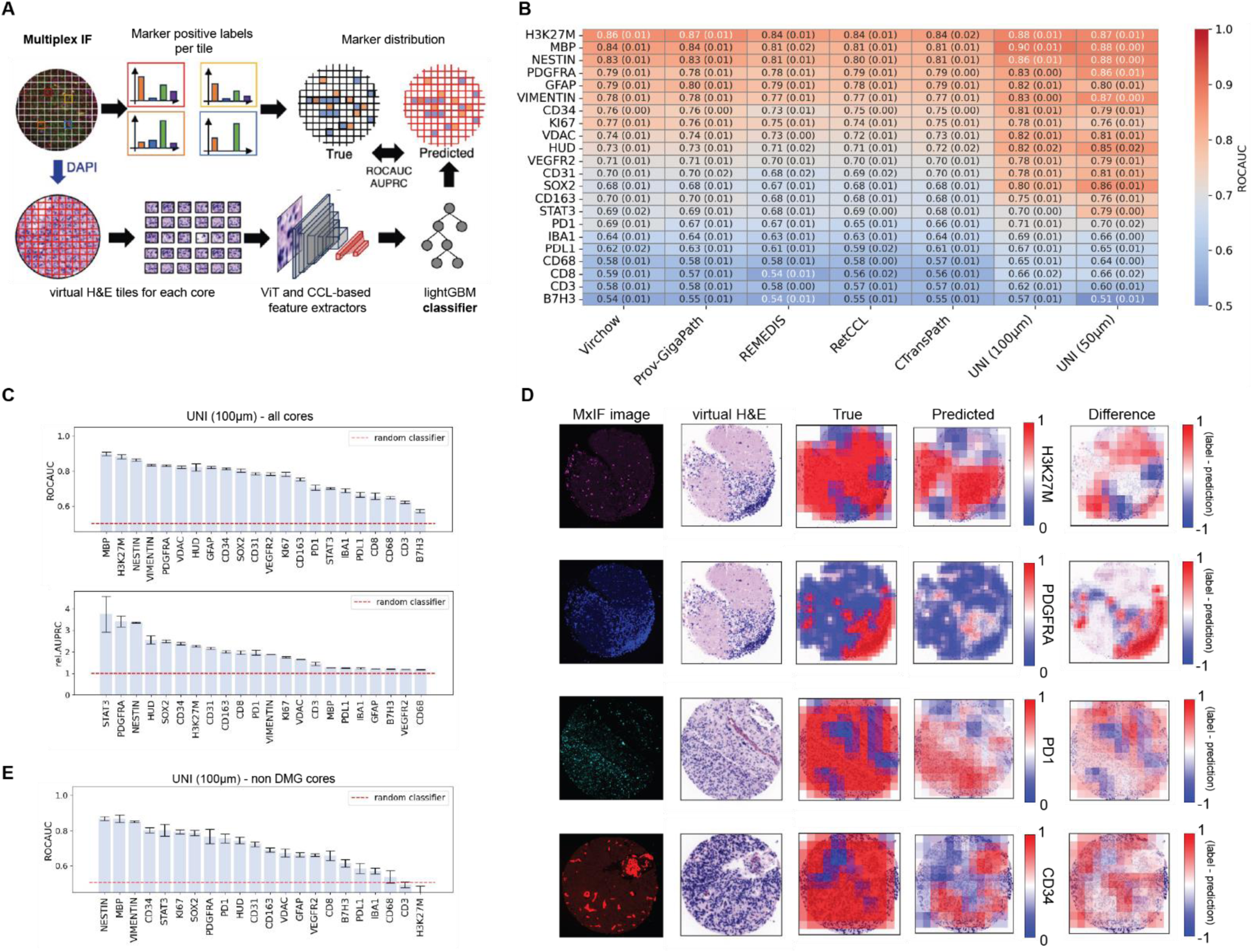
H&E-based prediction of cell markers in a digital pathology framework. (A) Workflow overview. Segmented TMA IF slides were cropped into tiles of 100 µm edge length and the presence/absence of positively stained cells for 22 IF markers was recorded. For each image tile a virtual H&E was created based on the DAPI signal and passed through a CCL-based feature extractor. For each marker a *lightGBM* classifier was trained on the extracted image features. We evaluate the performance in terms of ROCAUC and relative AUPRC between the true and predicted labels for all tiles within an independent test TMA. (B) Marker prediction performance overview across six state-of-the-art feature extractors, and two image tile resolutions (100µm and 50µm). Unless stated otherwise, image patch size was 100µm. ROCAUC is shown as mean values with standard deviations across five model instances train by nested cross-validation. All results refer to external test set data (TMA 2). The UNI vision transformer-based feature extractor was identified to outperform other approaches. We obtain competitive performance both at 100 and 50µm tile sizes. (C) Detailed results for UNI-based extracted features from 100µm tiles with downstream prediction performance. ROCAUC (top) and relative (rel.) AUPRC (normalized by the positive class prevalence) scores obtained on external test data (TMA 2) are shown. (D) Examples of spatial predictions of four example markers (top to bottom: H3K27M, PDGFRA, PD1, CD34). The IF signal and virtual H&E core images are shown together with the true label overlay (1 positive tile, 0 negative) and the predicted probability maps. Spatial difference maps between the true label and predicted class probability are given on the right. Results are shown for the best-performing feature extractor for the relevant marker, i.e. UNI-100µm, except for PDGFRA (UNI-50µm). (E) Subgroup prediction performance for primary tumor types different from DMG (on TMA 2).

## Discussion

Multiplex immunofluorescent imaging has emerged as a promising platform for mapping cellular neighborhoods across tumor and healthy tissue. The progress in reliable and reproducible multiplex imaging has resulted in mapping tumor microenvironment in non-small cell lung cancers [28], adult and pediatric low-grade gliomas [29], breast cancer [30], and several other malignancies. Here, we employed the MxIF multiplexed fluorescent microscopy method [31] and showed its ability for quantitative, subcellular, and single-cell mapping of a large cohort of brain tumor specimens. MxIF uses computational capturing of sequential images, aligned by nuclear counterstains, providing an opportunity for countless fluorescent immune probing [31, 32]. Similar studies have helped better understand IDH1 mutant pediatric CNS cancers by showing significant downregulation of vimentin, VEGFR2, nestin, and Ki67 in these cohort compared to IDH1 wildtype tumors [29].

We used MxIF to characterize the largest pediatric CNS tissue microarray published to date and focused on mapping immune engagement across DMGs in the context of metastasis and treatment modalities. We selected 44 antigens for our study, of which 33 showed reliable staining. This manuscript focused on 22 antigens that inform immune and brain cell sub-populations. We used these 22 biomarkers to map microenvironment across 816 punch cores representing 79 whole brains. Our TMA is unique and comprehensive given that it covers multiple anatomical regions within 79 pediatric whole brains. One major advantage of our study is the existence of matched frozen specimen where we performed validation studied (mRNA) across primary and metastatic sites. Indeed, our TMA multiplexed map for the 33 antigens, which passed the validation will be available digitally providing an opportunity for the scientific community to utilize this unique resource. Our observation of low CD3+ and CD8+T-cell abundant in DMG tumors is in line with publications which categorize DMGs as ‘cold’ tumors [3, 10, 33, 34].

However, our analyses indicated that DMG cold tumor immune microenvironment is not static. for example, we observed ramified to amoeboid morphological change of Iba1+ cells in primary DMG sites indicating activation of microglia cells (**Fig. 3A**) [35]. This morphological change was exacerbated in specimen from patients with ONC201, immunotherapy or combination therapies. Our data are alighted with the published preclinical data indicating immune activation and immune-cell infiltration following treatment with ONC201 [12, 36].

PDL1 is a tumor-associated immune suppressive marker which is reported to be minimally expressed in autopsied DIPG tumor specimen [15]. However, our data indicate that 37.7% (with some as high as 90%) of all cells in DMG primary sites to be PDL1+. However, only ∼15% of DMG cells were double positive for PDL1 and H3K27M+. This co-expression increased to up 88% in some DMG primary site; an observation that will need further investigations. Our observation of PDL1+-DMG cells interacting with PD1+ T-cells has been also observed in other tumors including large B-cell lymphomas [37, 38]. The exact nature of ligand-receptor interaction and their role in DMG microenvironment warrants further research.

Our observation of the relative lack of B7H3 was surprising. B7H3 is a transmembrane protein that is highly expressed by a number of tumor cells [39]. One of the first studies reporting B7H3 expression in DMGs used nine patient specimens collected at postmortem and reported 100% positivity across all nine specimens tested [40]. These studies resulted in B7H3-targeting immunotherapy clinical trials [9, 40–42]. Our data indicating the minimal B7H3 expression in H3K27M+ DMG cells, may be contributed by a number of factors including, antibody specificity, antigen masking, use of postmortem specimen or other factors. However, our data indicating the expression of B7H3 in healthy tissue is an important observation, highlighting the need for further studies to map clinically relevant antigens.

Finally, we also demonstrate the value of the generated data set in the context of computational pathology. Given the cost and labor-intensive nature of spatial proteomic analysis, computational models capable of providing biological insights from minimally processed samples, such as H&E stains [43, 44], hold tremendous potential, especially for rare tissue specimens. While a range of data-driven techniques have previously been applied for automated tumor (sub-) typing [45, 46], there is currently no method capable of predicting proteomic signatures at high spatial resolution (sub 100µm) for pediatric brain tumor samples. Furthermore, spatial proteomic AI predictions could streamline histopathology workflows and the retrospective analysis of rare archival H&E slides with unprecedented throughput and accuracy [24, 47]. Training such tools, requires large and diverse data ideally comprising multiple tumor types, locations and replicates jointly processed [48]. We present an AI pipeline that builds on large-scale pre-trained feature extractors and demonstrates strong potential to enable in-depth characterization of minimally processed samples. We compared six state-of-the-art preprocessing and prediction pipelines to provide proof of principle regarding the predictability of selected biomarkers from (virtual) H&E images in pediatric brain tumors. This approach was only feasible given the large number and diversity of samples processed as part of our TMA. We demonstrate that in particular the prediction of the H3K27M mutation is highly feasible, yielding competitive prediction performance and accurate spatial predictions of tumor boundaries and infiltrating regions. Our collective data indicate the opportunity to characterize additional pediatric CNS specimens for a comprehensive and detailed tumor TMA both in the context of tumor evolution (metastasis), and response to therapy. The lack of detection of selected biomarkers may in part be explained by the lower prevalence of cells positive for these markers, implying that a larger data set as well as specific training regimes could be employed to further investigate specific performance. H&E-stained tissue slides both from upfront biopsies, and post-mortem donations could now offer insight regarding the greater picture of the DMG and more generally pediatric brain tumor protein landscape ultimately informing clinical decision making.

## Methods

### Postmortem Donation processing

All autopsy samples were collected after written informed consent from the patient’s guardian before, at, or after death of the patient, as approved by the Institutional Review Board of Children’s National Hospital study entitled “Molecular Analysis of Pediatric Cancers” (#Pro00001339) or collected under the CNH-IRB approved exempt protocol “Biomarker Identification In Pediatric Brain Tumors” (#Pro00000747) or Swiss Ethical approval BASEC-Nr 2019-00615. Tumor tissue was obtained from patients at autopsy and stored in a deep freezer at −80°C after flash freezing in liquid nitrogen, fixed 10% neutral buffered formalin, or cryopreserved in DMSO supplemented freezing media. All patient identifiers were removed and re-assigned with research numerical identifiers for permanent storage within Children’s National Hospital and University Children’s Hospital Zurich. The coordination and procedure of collecting the samples was previously described in detail previously [49]. Human tissue samples stored in formalin were stored anywhere from 10 years to 2 weeks prior to undergoing the full embedding process.

### Embedding and production of tissue microarray (TMA)

Fixation, dehydration, and embedding in paraffin of the tissue (FFPE) was performed by standard protocols. A total of 623 samples were reviewed by Dr. Elisabeth Rushing, pathologist at University Children’s Hospital Zurich, and selected blocks were cut and stained for H&E. A total of 214 samples (242 FFPE blocks) from 69 patients and 10 non-CNS tumor patients as healthy control were included. Up to four brain sides were chosen per patient which were mostly brainstem, cerebellum, thalamus, and cortex. Brain sides were classified for primary tumor, metastatic, infiltration, and adjacent healthy area. Three punch cores (n=3) with a diameter of 0.6 mm were taken per tissue block based on H&E. Two TMAs were fabricated with a total of 918 punch cores with the TMA Grand Master Panoramic 250 Scanner 3DHistec. Control healthy tissue (non-CNS tumor patients) and ONC201 treated patients were included on both TMA blocks. TMA blocks had a dimension of 17 x 23 mm. For performing all described immunofluorescence staining 5 µm sections were cut.

Supplementary Table 1 summarizes number of patients, specimen, FFPE blocks, punch cores, markers used for each specific analysis (**Sup. Table 1**).

### Immunohistochemistry (IHC) staining

Immunohistochemistry (IHC) was performed on 2 µm sections to validate tissue punch cores on the TMAs after performing classical histology staining with H&E. Tumor markers H3K27M and H3K27me3 for DMG and Ki67 for other tumor types were used to validate dissemination sides. IHC was performed using the automated system BOND Fully Automated IHC Staining System (Leica Bond-III Processing Module | Biosystems Switzerland AG) with the dilution 1:100 protocol at University Hospital Zurich.

### Multiplexed immunofluorescence (MxIF) Cell DIVE™ imaging

MxIF imaging was conducted on 2 TMA slides of 5 µm thickness using Cell DIVE™ (Leica Microsystems, Issaquah, WA), which involves repeated cycles of staining, imaging, and signal inactivation (**Fig. 2A**). Imaging was performed by GE HealthCare using methodologies described previously [31]. In brief, a panel of 44 (including DAPI) antibodies was used to stain TMA slides after deparaffinization, rehydration, and a two-step, heat-mediated antigen retrieval using citrate (pH 6) and Tris (pH 9) process. Antigens were detected using either primary antibodies with fluorophore-labelled secondary antibodies or fluorescent dyes directly conjugated to primary antibodies. Prior to antibody staining, tissues were also stained with DAPI and imaged in all channels of interest to collect inherent tissue autofluorescence (AF). One field-of-view per core at 20X was collected each round using DAPI, Cy2, Cy3, Cy5, and Cy7 channels, followed by automated AF subtraction and registration with baseline DAPI. Pseudo-colored virtual H&E images were also generated using DAPI and AF signal. Tissue was stained with up to four antibodies per cycle for 1 hour at room temperature using the Leica Bond BOND-MAX autostainer, imaged, and then dye inactivated to remove signal. Cycling was repeated until all target antigens were probed **(Table 1)**.

The typical antibody characterization workflow is a standardized process that consists of screening at least three commercial antibody clones per target for sensitivity and sensitivity, evaluation of the epitope for susceptibility to the dye inactivation workflow, and concordance of antibody performance upon dye conjugation [50].

### TMA analysis using QuPath

QuPath (v.0.3.2) was used to visualize the TMA MxIF images, as well as to perform cell segmentation, feature extraction (mean intensity per marker, x, y centroid coordinate) per single cell, and for counting the relative proportion of positive cells for a given marker. Cell segmentation was performed using the “Cell detection” tool implemented in QuPath across all cores of the TMA using the DAPI channel. Default segmentation parameters were used: background radius 8 µm, minimum area 10 µm^2^, maximum area 400 µm^2^, cell expansion of 5 µm. The intensity parameter threshold for DAPI was chosen between 250 to 1000 depending on the core. Due to the age of the postmortem tissue and different fixation times in formaldehyde, the background signal of markers varied between patient brain areas and cores. Single and composite classifiers were generated to set thresholds depending on background intensity per core. We counted positive cells for a given marker and threshold using the QuPath “TMA measurement” function and merged the counts across the three cores per patient (n=3) together. Positive cell numbers were put into relative ratio with the total number of cells in the core and triplicates (3 punch cores per sites) were combined (n=3). Specific markers, B7H3 and PDL1, were put additionally into relative ratio with DMG tumor cells (H3K27M+ cells) to investigate double positive cells. GraphPad Prism (v.10.0.2 (232) was used for visualization and to perform statistical tests using the Kruskal-Wallis (H-Test) and Mann-Whitney (U-Test) test, Unpaired t-test, One-way ANOVA and 2-way ANOVA, Šídák’s multiple comparisons test as appropriate and indicated in the figure legends.

### Single-cell spatial analysis

Raw TIFF images and cell segmentations boundaries were summarized into a single cell (n = 762’988 cells) marker expression matrix using QuPath “Detection measurements” function by calculating the mean marker intensity within either the nucleus, cell (expanded boundary), or cytoplasm (cell minus nucleus) boundaries. The quality of the staining for each channel was manually checked and the mean signal (for n = 22 markers) within the following segmentation boundaries was considered for downstream analysis:

We removed cores that displayed high levels of background signal, where thresholds could not be set, or no signal for DAPI. For each patient (1 to 3 cores), cells were independently annotated (i.e. phenotyping) before merging the data. For each channel, the raw data was re-scaled using an inverse hyperbolic sine function and a cofactor of 5. For patients with more than one core, batch/core correction was performed using Harmony [51] on the scaled data. Following batch correction, cells sharing similar marker expression profiles were grouped together by Louvain clustering using k = 50 nearest neighbors. Cell types were manually assigned based on relative marker expression values and cross-checking raw signal on MxIF images. The annotated data was then merged into a SpatialExperiment object [52] for UMAP and spatial visualization using in R (v.4.3.1) and the imcRtools (v.3.19) [53] package.

### Biomarker and cell type prediction from H&E image tiles

Coordinates (x-y) of cells on MxIF images were extracted using QuPath software leading to a co-localization matrix indicating for each cell the relevant IF intensity and location. This enables co-localization with a cell’s pseudo-colored virtual H&E image and IF profile. We threshold the IF for all 22 markers as described in section “TMA analysis using QuPath” to obtain binary labels for each cell.

Virtual H&E images of all cores were sub divided into partially (up to 75%) overlapping quadratic tiles of 50, or 100 µm edge length obtained by shifting a selection window first horizontally, then vertically by half the edge length. Any tile comprising at least one cell (5 cells for 100µm tiles) was included for predictions. A tile was labeled positive for each of the 22 markers if at least one positively stained cell was present, leading to 22 separate label vectors. We use TMA1 data for model training and separate these into 5-fold cross-validation fold (i.e. 20% internal test data) stratified at patient level based on DMG tumor presence/absence.

Six pretrained frameworks were benchmarked for feature extraction from H&E images. These included two architectures published in 2023 based on Res-Nets (RetCCL [47], REMEDIS [23]), CTransPath [26] as a popular approach based on a Swin Transformer, as well as three vision transformer-based architectures published in 2024: Prov-GigaPath [27], UNI [24], and the Virchow et al. model [25]. We summarize key characteristics for these approaches below:

| Feature extractor | Architecture | Training Data | Training regime | Features extracted |
| --- | --- | --- | --- | --- |
| RetCCL | Res-Net-50 | TCGA & PAIP | Clustering based contrastive learning | 2048 |
| REMEDIS | Res-Net-152 | TCGA | Contrastive learning (SimCLR) | 4096 |
| CTransPath | Swin transformer | TCGA & PAIP | Contrastive learning (MoCoV3) | 768 |
| Prov-GigaPath | Vision transformer (GigaPath) for gigapixel pathology slides | 171'000 WSI from Providence health network | Mask image modeling and self-distillation (DINOv2 and LongNet) | 1536 |
| UNI | Vision transformer | Mass-100k (MGH & BWH) | Mask image modeling and self-distillation (DINOv2) | 1024 |
| Virchow | Vision Transformer | 1,488,550 WSIs from MSKCC | Mask image modeling and self-distillation (DINOv2) | 2560 |

Features were extracted from virtual H&E image tiles in *.png* format. Depending on the feature extractor, 768 to 4096 features were used. These features were used to train a binary prediction classifier to identify within each patch the presence/absence of marker positive cells: All features were standard-scaled using training set means and standard deviations. Features of TMA 2 were scaled accordingly based on the relevant cross-validation fold. We performed an image-feature-based classification task using lightGBM, a popular tree-based ensemble model framework available in the Python *lightgbm* (version 4.3.0) library. A separate model was trained for each of the n = 22 markers exclusively on TMA 1 data. Hyperparameter optimization was performed using *Gridsearch* in a nested 3-fold cross-validation to optimize the learning rate, number of leaves, and regularization. In total ∼47k/187k 100/50µm tiles originating from 420 cores were used for model training.

In addition to validation in the left-out TMA 1 test sets, TMA 2 (405 cores, 45k or 174k; 100µm or 50µm tiles) served as an independent test data set for the evaluation of prediction performance. We evaluate performance using the area under the receiver operator characteristic (ROCAUC) as well as the area under the precision-recall curve (AUPRC) to provide a metric more suitable for imbalanced data. As AUPRC of a random classifier is dictated by the positive class prevalence, we normalize these values by the positive class prevalence. Additionally, we visualize true and predicted image patch labels overlaid to the original pseudo-colored virtual H&E image. All models were implemented in Python (v. 3.11.6) based on the *scikit-learn (1.3.2)* and *lightgbm* (4.3.0) libraries. All relevant codes are available on Github. (https://gitlab.ethz.ch/BMDSlab/publications/dmg_tma_public.git)

### Biomarker validation in human plasma using multiplex ELISA technology Mesoscale Discovery (MSD)

To validate biomarkers found in the MxIF staining, human patients’ plasma samples collected at three different time points in their treatment and treated with ONC201 (n=3) and ONC201 naïve (n=3) as well as non-tumor patients were analyzed using Mesoscale Discovery (MSD) U-Plex plate systems. Biomarker panel was chosen from U-PLEX Custom Immuno-Oncology Grp 1 (hu) Assays, (MSD, K151AEM-1). The following panel was used in this study: CD27, CD20, B7H3 and PD1. The experiment was performed following manufacturers protocol. In brief, plasma samples were thawed on ice and used as a 1:5 dilution. Plates were coated over night with linker and antibody solution, followed by washing step and stored overnight at 4°C. Calibrator and samples were added to the plate and incubated for 2h at RT followed by wash steps and adding of detection antibody and incubation for 1 hr. Statistical analysis was done using GraphPad Prism (v.10.0.2 (232)) and 2-way ANOVA test.

### FACS analysis of viably frozen patient tissue

For FACS analysis, postmortem tissue was processed directly after autopsy by mincing tissue in culture medium and finally freezing down in culture medium supplemented with DMSO. 3 patients and two brain sides, including primary tumor and adjacent tumor side was used to study immune cell composition. Tissue samples were processed for FACS accordingly. In brief, by vials were thawed and washed in TSM compete media through centrifugation, 300xg, 5 minutes. Minced tissue pellets were triturated gently and homogenized in TSM complete media using a 100 μm strainer and the plunger from a 1 mL syringe. Suspensions were left to settle for 3-5 minutes and supernatants were transferred to a clean tube. Samples were centrifuged and cell pellets were again filtered, resuspended in TSM complete media, and transferred to a T25 low attach flask for incubation at 37°C o/n*. The day after, cell suspensions were collected for flow cytometry.

The following antibody panels were used:

Panel 1: CD4/8/25/27/PD1/TIGIT, 28/39

Panel 2: CD45/HLADR/123/14/16/33/36/SLAN

Panel 3: CD4/8/45RA/62L/CCR4/27/CXCR3/CCR6

Panel 4: CD3/8/TIM3/38/56/PD1/TIGIT/37

Cells were acquired in a FACSCanto-II (BD Biosciences). Data were analyzed with Infinicyt 2.0 software (Cytognos) and represented using GraphPad Prism.

### Validation of immune markers on mice tissue treated with ONC201 by IHC

26C-7 cells were implanted orthotopically into the pons, following the next coordinates with lambda as the reference point (Y: 1.5mm, X: 0.8mm, Z: 5 mm) in C57BL6/J mice. The cell line was kindly provided by Timothy N. Phoenix (Cincinnati Children’s Hospital Medical Center, Cincinnati) and are intrauterine electroporation (IUE)-derived cells, containing dominant-negative p53 (DN-p53), PDGFRAD842V and H3.1K27M/ACVR1G328V. 250,000 cells were implanted at day 0. After 19 days ONC201 mice were treated with 125 mg/kg dosage once per week, afterwards sacrificed at similar timepoints after receiving treatment 5-6 times with vehicle or ONC201 administration. In this case, three tumor sections of each mouse (3 mice in total = 9 sections in total) per condition were analyzed in the graphs.

For IHC staining, paraffin sections (2µm thick) were cut, dewaxed and hydrated. Antigen retrieval was performed in a Pascal pressure chamber (S2800, Dako) for 30 min at 95 °C in 10 mM Tris-1 mM EDTA solution pH 9. Slides were allowed to cool for 20 min, then endogenous peroxidase was blocked with 3% H_2_O_2_ in deionized water for 12 min. After washing in TBS-0,05% Tween 20 (TBS-T), sections were incubated overnight at 4°C with primary antibodies. After washing in TBS-T, goat anti-rabbit labelled polymer EnVisionTM+ System (K400311-2, Agilent) was applied for 30 min at room temperature and peroxidase activity was revealed using DAB+ (K346811-2, Agilent). Finally, sections were lightly counterstained with Harris hematoxylin, dehydrated, and cover slipped with Eukitt (Labolan, 28500). This was done with the kindly help and support of the Morphology Core Facility (Universidad de Navarra, Centro de Investigación Médica Aplicada-CIMA-). The primary antibodies used were the following: **CD3:** Neomarkers, RM9107; 1:300, **CD8:** Cell Signaling, 98941; 1:400, **F4/80:** Cell signaling, 70076; 1:2000, **Iba1:** Wako, 019-19741; 1:4000

### RNA Sequencing and analysis

Fresh frozen (5 mg) brain tissue samples were processed for RNA isolation, including primary tumor (n = 59) and metastatic sites (n = 28) from 62 patients. RNA was extracted by using the RNeasy Plus Micro Kit (Qiagen, 74034). RNA concentration was measured by Nanodrop and Qubit. A minimum of 200 ng RNA was used for the library preparation using RNA-seq - TruSeq mRNA-seq library and followed by Illumina NovaSeq 6000, 50PE sequencing to 25M clusters per library.

RNA-seq data were obtained as FASTQ files from the total of 87 samples and were processed using the nf-core RNA-sequence pipeline (v 3.14.0) [54] with default parameters. Briefly, quality control of raw reads was assessed using FastQC [55], adaptor sequences and low-quality reads were trimmed using Trim Galore [56], and then aligned to the human reference genome (GRCh38) using STAR aligner (version 2.7.3a) [57]. Gene-level quantification was performed using Salmon (version 1.4.0) [58], resulting in a gene count matrix used for downstream differential expression analysis using the DESeq2 package (v. 1.32.0) [59]. Genes with zero counts across all samples were removed to reduce noise and enhance statistical power. One sample was identified as low quality and was excluded from downstream analyses.

To explore sample-to-sample relationships and visualization, we performed Uniform Manifold and Projection (UMAP) and computed pairwise Euclidean distances on VST-transformed data and using the expression levels of the top 1000 most variable genes. The VST normalization was used to transform count data while stabilizing the variance across the mean. UMAP dimensionality reduction was conducted using the umap package with default parameters. Two groups of samples were identified from the UMAP (group A and group B). The distance matrix was calculated across samples using the dist function and hierarchically clustered using the pheatmap package [60].

Differential expression analysis was performed across DMG primary and metastasis samples, ONC201 treated vs non-treated DMG primary samples, and between group A and group B in DMG samples, by subsetting the corresponding data and specifying contrasts of interest. Significantly differentially expressed genes were identified with an adjusted p-value cutoff of 0.05 (Benjamini-Hockberg correction to control for false discovery rate) and vizualized as a volcano plot. Immune-related genes were identified based on Gene Ontology (GO) terms associated with immune response, specifically GO:0006955 (immune response), GO:0002250 (adaptive immune response), and GO:0002376 (immune system process). Gene annotations were retrieved using biomaRt [61], and immune-related protein-coding genes were selected for downstream analyses.

Functional enrichment analysis was performed using the g:Profiler tool and gprofiler2 package [62]. Upregulated and downregulated genes were defined as those with log2 fold changes greater than 1 or less than –1, respectively. Adjusted p-values were obtained using the g:SCS method, which controls for multiple testing by considering the structure of the Gene Ontology.

Significance was considered for adjusted p-values less than 0.05. Plots were generated using the ggplot2 and pheatmap packages.

The RNAseq data and single-cell proteomics atlas will be deposited on GEO after the manuscript is accepted.

## Acknowledgment

We would like to thank patients and their families for donating specimen funds for our research. We would like to thank the Rising Tide Foundation for their generous financial support of this work, LilaBean Foundation, The Swifty Foundation, Gift From a Child (GFAC) Initiative, Swiss to Cure DIPG, Yuvaan Tiwari Foundation, Basel Research Center for Child Health.

## Authors Contributions

**S.L., A.J.D.M., S.B.** contributed conceptualization, Data curation, Formal analysis, investigation, visualisation, writing-original draft, writing-review & editing.

**J.N.** contributed conceptualization, Data curation, Formal analysis, investigation, visualisation, writing-original draft, writing-review & editing, Supervision, Funding Acquisition.

**A.P.** contributed conceptualization, Formal analysis, investigation, writing-original draft, writing-review&editing.

**E.M., E.J.R., C.S., F.G., S.D., F.P., N.C.L., D.N.** contributed to investigation.

**J.B.** contributed to investigation, Formal analysis, visualization.

**A.E., L.K., D.M., M.B.** contributed resources.

**L.M.** contributed to visualization.

**M.D.D., M.G., A.S.G.S., S.M., R.P., M.M.A.** contributed to writing-review & editing.

## Literature

1. Cunningham, R.M., M.A. Walton, and P.M. Carter, The Major Causes of Death in Children and Adolescents in the United States. N Engl J Med, 2018. 379(25): p. 2468–2475.

2. Noon, A. and S. Galban, Therapeutic avenues for targeting treatment challenges of diffuse midline gliomas. Neoplasia, 2023. 40: p. 100899.

3. Persson, M.L., et al., The intrinsic and microenvironmental features of diffuse midline glioma: Implications for the development of effective immunotherapeutic treatment strategies. Neuro Oncol, 2022. 24(9): p. 1408–1422.

4. Louis, D.N., et al., The 2021 WHO Classification of Tumors of the Central Nervous System: a summary. Neuro Oncol, 2021. 23(8): p. 1231-1251.

5. Vallero, S.G., et al., Pediatric diffuse midline glioma H3K27-altered: A complex clinical and biological landscape behind a neatly defined tumor type. Front Oncol, 2022. 12: p. 1082062.

6. Schwartzentruber, J., et al., Driver mutations in histone H3.3 and chromatin remodelling genes in paediatric glioblastoma. Nature, 2012. 482(7384): p. 226-31.

7. DeNunzio, N.J. and T.I. Yock, Modern Radiotherapy for Pediatric Brain Tumors. Cancers (Basel), 2020. 12(6).

8. Gallitto, M., et al., Role of Radiation Therapy in the Management of Diffuse Intrinsic Pontine Glioma: A Systematic Review. Adv Radiat Oncol, 2019. 4(3): p. 520–531.

9. Lieberman, N.A.P., et al., Characterization of the immune microenvironment of diffuse intrinsic pontine glioma: implications for development of immunotherapy. Neuro Oncol, 2019. 21(1): p. 83–94.

10. Andrade, A.F., et al., Immune landscape of oncohistone-mutant gliomas reveals diverse myeloid populations and tumor-promoting function. Nat Commun, 2024. 15(1): p. 7769.

11. Pachocki, C.J. and E.M. Hol, Current perspectives on diffuse midline glioma and a different role for the immune microenvironment compared to glioblastoma. J Neuroinflammation, 2022. 19(1): p. 276.

12. Jackson, E.R., et al., ONC201 in combination with paxalisib for the treatment of H3K27-altered diffuse midline glioma. Cancer Res, 2023. 83(14): p. 2421–37.

13. Jackson, E.R., et al., A review of current therapeutics targeting the mitochondrial protease ClpP in diffuse midline glioma, H3 K27-altered. Neuro Oncol, 2024. 26(Supplement_2): p. S136-S154.

14. Gallego Perez-Larraya, J., et al., Oncolytic DNX-2401 Virus for Pediatric Diffuse Intrinsic Pontine Glioma. N Engl J Med, 2022. 386(26): p. 2471-2481.

15. Chen, Y., et al., Immune Microenvironment and Immunotherapies for Diffuse Intrinsic Pontine Glioma. Cancers (Basel), 2023. 15(3).

16. Vitanza, N.A., et al., Intraventricular B7-H3 CAR T Cells for Diffuse Intrinsic Pontine Glioma: Preliminary First-in-Human Bioactivity and Safety. Cancer Discov, 2023. 13(1): p. 114–131.

17. Harms, P.W., et al., Multiplex Immunohistochemistry and Immunofluorescence: A Practical Update for Pathologists. Mod Pathol, 2023. 36(7): p. 100197.

18. Duhamel, M., et al., Spatial analysis of the glioblastoma proteome reveals specific molecular signatures and markers of survival. Nat Commun, 2022. 13(1): p. 6665.

19. Piwecka, M., N. Rajewsky, and A. Rybak-Wolf, Single-cell and spatial transcriptomics: deciphering brain complexity in health and disease. Nat Rev Neurol, 2023. 19(6): p. 346–362.

20. Jin, L., et al., Artificial intelligence neuropathologist for glioma classification using deep learning on hematoxylin and eosin stained slide images and molecular markers. Neuro Oncol, 2021. 23(1): p. 44–52.

21. Gadermayr, M. and M. Tschuchnig, Multiple instance learning for digital pathology: A review of the state-of-the-art, limitations & future potential. Comput Med Imaging Graph, 2024. 112: p. 102337.

22. Van Gool, S.W., et al., Addition of Multimodal Immunotherapy to Combination Treatment Strategies for Children with DIPG: A Single Institution Experience. Medicines (Basel), 2020. 7(5).

23. Azizi, S., et al., Robust and data-efficient generalization of self-supervised machine learning for diagnostic imaging. Nat Biomed Eng, 2023. 7(6): p. 756–779.

24. Chen, R.J., et al., Towards a general-purpose foundation model for computational pathology. Nat Med, 2024. 30(3): p. 850–862.

25. Vorontsov, E., et al., A foundation model for clinical-grade computational pathology and rare cancers detection. Nat Med, 2024. 30(10): p. 2924–2935.

26. Wang, X., et al., Transformer-based unsupervised contrastive learning for histopathological image classification. Med Image Anal, 2022. 81: p. 102559.

27. Xu, H., et al., A whole-slide foundation model for digital pathology from real-world data. Nature, 2024. 630(8015): p. 181-188.

28. Parra, E.R., et al., Immuno-profiling and cellular spatial analysis using five immune oncology multiplex immunofluorescence panels for paraffin tumor tissue. Sci Rep, 2021. 11(1): p. 8511.

29. Berens, M.E., et al., Multiscale, multimodal analysis of tumor heterogeneity in IDH1 mutant vs wild-type diffuse gliomas. PLoS One, 2019. 14(12): p. e0219724.

30. Clarke, G.M., et al., A novel, automated technology for multiplex biomarker imaging and application to breast cancer. Histopathology, 2014. 64(2): p. 242–55.

31. Gerdes, M.J., et al., Highly multiplexed single-cell analysis of formalin-fixed, paraffin-embedded cancer tissue. Proc Natl Acad Sci U S A, 2013. 110(29): p. 11982–7.

32. Wang, J., et al., Multiplexed immunofluorescence identifies high stromal CD68(+)PD-L1(+) macrophages as a predictor of improved survival in triple negative breast cancer. Sci Rep, 2021. 11(1): p. 21608.

33. Lin, G.L., et al., Non-inflammatory tumor microenvironment of diffuse intrinsic pontine glioma. Acta Neuropathol Commun, 2018. 6(1): p. 51.

34. Hanahan, D. and M. Monje, Cancer hallmarks intersect with neuroscience in the tumor microenvironment. Cancer Cell, 2023. 41(3): p. 573–580.

35. Vidal-Itriago, A., et al., Microglia morphophysiological diversity and its implications for the CNS. Front Immunol, 2022. 13: p. 997786.

36. de la Nava, D., et al., The oncolytic adenovirus Delta-24-RGD in combination with ONC201 induces a potent antitumor response in pediatric high-grade and diffuse midline glioma models. Neuro Oncol, 2024.

37. Li, L., et al., PD-1/PD-L1 expression and interaction by automated quantitative immunofluorescent analysis show adverse prognostic impact in patients with diffuse large B-cell lymphoma having T-cell infiltration: a study from the International DLBCL Consortium Program. Mod Pathol, 2019. 32(6): p. 741–754.

38. Zitvogel, L. and G. Kroemer, Targeting PD-1/PD-L1 interactions for cancer immunotherapy. Oncoimmunology, 2012. 1(8): p. 1223–1225.

39. Feustel, K., J. Martin, and G.S. Falchook, B7-H3 Inhibitors in Oncology Clinical Trials: A Review. J Immunother Precis Oncol, 2024. 7(1): p. 53–66.

40. Zhou, Z., et al., B7-H3, a potential therapeutic target, is expressed in diffuse intrinsic pontine glioma. J Neurooncol, 2013. 111(3): p. 257–64.

41. Gregorio, A., et al., Small round blue cell tumours: diagnostic and prognostic usefulness of the expression of B7-H3 surface molecule. Histopathology, 2008. 53(1): p. 73–80.

42. Majzner, R.G., et al., CAR T Cells Targeting B7-H3, a Pan-Cancer Antigen, Demonstrate Potent Preclinical Activity Against Pediatric Solid Tumors and Brain Tumors. Clin Cancer Res, 2019. 25(8): p. 2560–2574.

43. Marini, N., et al., Data-driven color augmentation for H&E stained images in computational pathology. J Pathol Inform, 2023. 14: p. 100183.

44. Lu, M.Y., et al., Data-efficient and weakly supervised computational pathology on whole-slide images. Nat Biomed Eng, 2021. 5(6): p. 555–570.

45. Lee, R.Y., et al., The promise and challenge of spatial omics in dissecting tumour microenvironment and the role of AI. Front Oncol, 2023. 13: p. 1172314.

46. Cui, M. and D.Y. Zhang, Artificial intelligence and computational pathology. Lab Invest, 2021. 101(4): p. 412–422.

47. Wang, X., et al., RetCCL: Clustering-guided contrastive learning for whole-slide image retrieval. Med Image Anal, 2023. 83: p. 102645.

48. Kumar, N., et al., A Dataset and a Technique for Generalized Nuclear Segmentation for Computational Pathology. IEEE Trans Med Imaging, 2017. 36(7): p. 1550–1560.

49. Kambhampati, M., et al., Harmonization of postmortem donations for pediatric brain tumors and molecular characterization of diffuse midline gliomas. Sci Rep, 2020. 10(1): p. 10954.

50. Chadwick, C., et al., Pathobiology of Candida auris infection analyzed by multiplexed imaging and single cell analysis. PLoS One, 2024. 19(1): p. e0293011.

51. Korsunsky, I., et al., Fast, sensitive and accurate integration of single-cell data with Harmony. Nat Methods, 2019. 16(12): p. 1289–1296.

52. Righelli, D., et al., SpatialExperiment: infrastructure for spatially-resolved transcriptomics data in R using Bioconductor. Bioinformatics, 2022. 38(11): p. 3128–3131.

53. Windhager, J., et al., An end-to-end workflow for multiplexed image processing and analysis. Nat Protoc, 2023. 18(11): p. 3565–3613.

54. Ewels, P.A., et al., The nf-core framework for community-curated bioinformatics pipelines. Nat Biotechnol, 2020. 38(3): p. 276–278.

55. Andrews, S. FastQC: A Quality Control Tool for High Throughput Sequence Data. 2010; Available from: https://github.com/s-andrews/FastQC.

56. Krueger, F. Trim Galore! A wrapper tool around Cutadapt and FastQC to consistently apply quality and adapter trimming to FastQ files. 2015; Available from: https://github.com/FelixKrueger/TrimGalore.

57. Dobin, A., et al., STAR: ultrafast universal RNA-seq aligner. Bioinformatics, 2013. 29(1): p. 15–21.

58. Patro, R., et al., Salmon provides fast and bias-aware quantification of transcript expression. Nat Methods, 2017. 14(4): p. 417–419.

59. Love, M.I., W. Huber, and S. Anders, Moderated estimation of fold change and dispersion for RNA-seq data with DESeq2. Genome Biol, 2014. 15(12): p. 550.

60. Kolde, R. pheatmap: Pretty Heatmaps. R package version 1.0.12. 2018; Available from: https://github.com/raivokolde/pheatmap.

61. Smedley, D., et al., BioMart--biological queries made easy. BMC Genomics, 2009. 10: p. 22.

62. Reimand, J., et al., g:Profiler-a web server for functional interpretation of gene lists (2016 update). Nucleic Acids Res, 2016. 44(W1): p. W83–9.

