## Supplemental material for "Mapping Tumor Microenvironment and Treatment Response of Diffuse Midline Glioma Using Multiplexed Immunofluorescence and AI Models"

| Type | Immunohistochemistry (IHC) (TMA) | Immunofluorescence (IF) (TMA) | RNAseq |
|---|---|---|---|
| Type of specimen | FFPE sections | FFPE sections | Frozen tissue |
| # patients | 79 (DMG 49) | 76 (DMG: 47, OS: 37) | 62 (DMG 47) |
| # Specimen | 224 (DMG 157) | 224 (DMG 157) | 87 (DMG 68) |
| # FFPE blocks | 242 (DMG 167) | 242 (DMG 167) | - |
| # punch cores | 906 (DMG 627) | 816 (DMG 558) | - |
| # markers stained | 4 (+ H&E) | 44 | - |
| # markers passed | 4 (+ H&E) | 33 | - |
| # markers reported | 3 (+ H&E) | 22 | - |

**Sup. Table 1**: Overview of cohort, sample size, number of markers taken for the listed analysis IHC, IF, RNA sequencing.

A

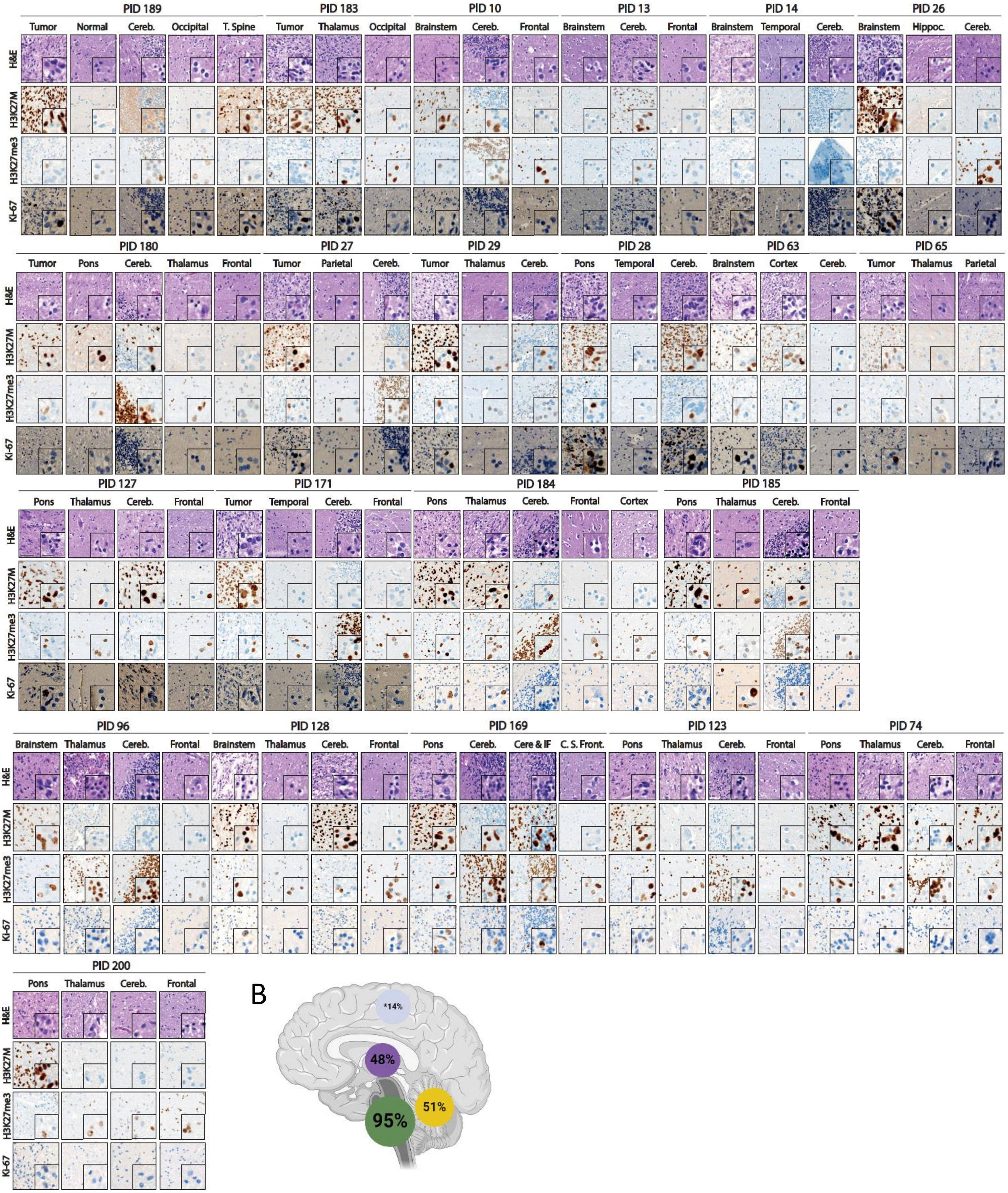

PID 189
Tumor
Normal
Cereb.
Occipital
T. Spine
PID 183
Tumor
Thalamus
Occipital
PID 10
Brainstem
Cereb.
Frontal
PID 13
Brainstem
Cereb.
Frontal
PID 14
Brainstem
Temporal
Cereb.
PID 26
Brainstem
Hippoc.
Cereb.
H&E
H3K27M
H3K27me3
Ki-67
PID 180
Tumor
Pons
Cereb.
Thalamus
Frontal
PID 27
Tumor
Parietal
Cereb.
PID 29
Tumor
Thalamus
Cereb.
PID 28
Pons
Temporal
Cereb.
PID 63
Brainstem
Cortex
Cereb.
PID 65
Tumor
Thalamus
Parietal
PID 127
Pons
Thalamus
Cereb.
Frontal
PID 171
Tumor
Temporal
Cereb.
Frontal
PID 184
Pons
Thalamus
Cereb.
Frontal
Cortex
PID 185
Pons
Thalamus
Cereb.
Frontal
PID 96
Brainstem
Thalamus
Cereb.
Frontal
PID 128
Brainstem
Thalamus
Cereb.
Frontal
PID 169
Pons
Cereb.
Cere & IF
C. S. Front.
PID 123
Pons
Thalamus
Cereb.
Frontal
PID 74
Pons
Thalamus
Cereb.
Frontal
PID 200
Pons
Thalamus
Cereb.
Frontal
B
*14%
48%
51%
95%

**Sup. Figure 1: Representative image of 28 DMG patients included on the TMA.**

(A) H&E, H3K27M, H3K27me3 and Ki67 staining of 28 representative DMG patients, including tissue cores from primary, metastatic and adjacent healthy brain sites to assess tumor and non-tumor regions. H3K27M and H3K27me3 loss confirmed tumor sites of DMG patients, whereas Ki67 showed highly proliferating sites.

(B) To assess primary, metastatic and adjacent healthy sites, H3K27M positive scored sites, done by a neuropathologist, across the DMG cohort (n = 49) were analyzed and revealed 95% of patients with positivity in brainstem, 51% in cerebellum, 48% in thalamus, 14% in cortical regions (Created with BioRender.com).

| Target | Clone | Vendor | Catalog |
|---|---|---|---|
| **14-3-3-sigma** | 1433S01 | Thermo Fisher | MS-1185-PABX |
| **Akt** | C67E7 | Cell Signaling | 4691S |
| **B7-H3** | MIH42 | BioLegend | 351007 |
| **Beclin** | polyclonal | Millipore Sigma | PRS3613 |
| **beta-actin** | 13E5 | Cell Signaling | 4970 |
| **CD163** | EDHu-1 | BioRad | MCA1853 |
| **CD20** | EP459Y | Abcam | ab198943 |
| **CD274 (PDL1)** | 28-8 | Abcam | ab205921 |
| **CD3** | F7.2.38 | Agilent | M725429 |
| **CD31** | 89C2 | Cell Signaling | 3528BF |
| **CD34** | QBEnd/10 | Leica | END-L-CE |
| **CD4** | EPR6855 | Abcam | ab181724 |
| **CD44** | 156-3C11 | Cell Signaling | 3570B |
| **CD44 V6** | VFF-7 | Thermo Fisher | BMS116 |
| **CD45** | 2B11 + PD7/26 | Agilent | M0701 |
| **CD47** | polyclonal | R&D | AF4670 |
| **CD68** | KP1 | Thermo Fisher | MS-397-PABX |
| **CD8** | C8/144B | Agilent | M7103 |
| **CD86** | BU63 | Santa Cruz | sc-19617 |
| **FOXG1** | EPR18987 | Abcam | ab270103 |
| **FOXO1** | C29H4 | Cell Signaling | 2880BF |
| **FOXO3a** | D19A7 | Cell Signaling | 14592 |
| **FOXP3** | 206D | BioLegend | 320114 |
| **GFAP** | GA5 | Thermo Fisher | 53 9892 82 |
| **H3K27M** | D3B5T | Cell Signaling | 85023 |
| **H3K27me3** | C36B11 | Cell Signaling | 5499 |
| **Hif1alpha** | EP1215Y | Abcam | ab51608 |
| **HuD** | E-1 | Santa Cruz | sc-28299 |
| **Iba1** | polyclonal | Wako chemical | 019-19741 |
| **Ki67** | SP6 | Abcam | ab197547 |
| **MBP** | 12 | Abcam | ab7349 |
| **NaKATPase** | EP1845Y | Abcam | ab197713 |
| **Nestin** | 10C2 | BioLegend | 656810 |
| **NeuN** | A60 | Millipore Sigma | MAB377 |
| **PD1** | EPR4877(2) | Abcam | ab137132 |
| **PDGFRa** | polyclonal | R&D | AF307 |
| **Erk1_2_pT202_pY204** | 20G11 | Cell Signaling | 4376BF |
| **mTor_pS2448** | 49F9 | Cell Signaling | 2976BF |
| **SOX2** | D6D9 | Cell Signaling | 5067S |
| **Stat3** | 124H6 | Cell Signaling | 9139B |
| **VDAC** | 20B12AF2 | Abcam | ab14734 |
| **VEGFR2** | 55B11 | Cell Signaling | 2479 |
| **Vimentin** | D21H3 | Cell Signaling | 5741BF |
| **DAPI nucleus stain** | | | |

**Sup. Table 2**: **44 antibodies used in this study with clone information and catalogue number.**

Antibodies were selected to cover brain specific healthy cells (Neurons, Astrocytes, Oligodendrocytes), DMG tumor cells, immune cells (lymphocytes, myeloid and immune suppressive phenotypes), proliferative cells, stem cells and endothelial cells. Further some metabolic markers and tumor specific markers were included.

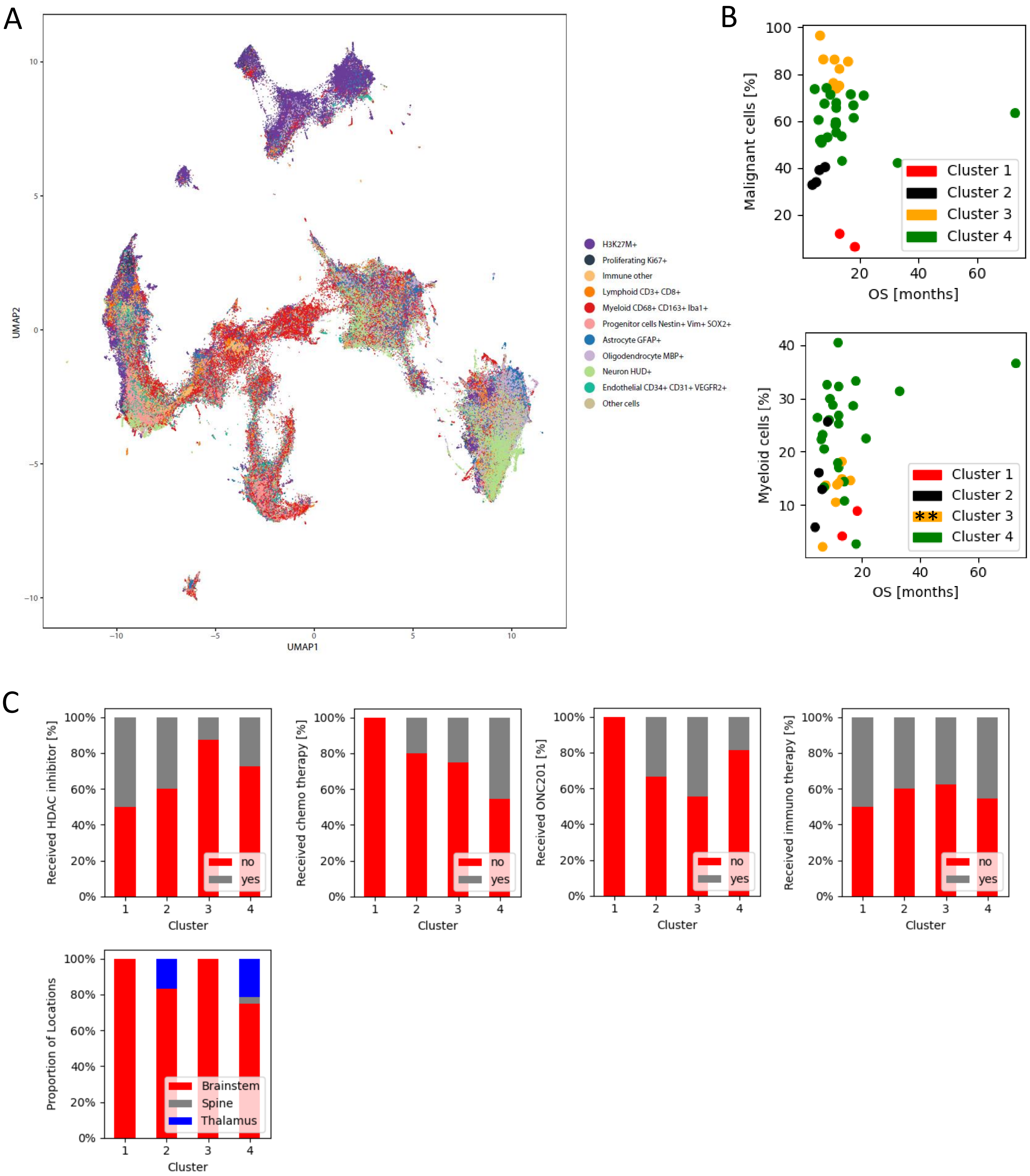
A
B
C
UMAP1
UMAP2
H3K27M+
Proliferating Ki67+
Immune other
Lymphoid CD3+ CD8+
Myeloid CD68+ CD163+ Iba1+
Progenitor cells Nestin+ Vim+ SOX2+
Astrocyte GFAP+
Oligodendrocyte MBP+
Neuron HUD+
Endothelial CD34+ CD31+ VEGFR2+
Other cells
Malignant cells [%]
Myeloid cells [%]
OS [months]
Cluster 1
Cluster 2
Cluster 3
Cluster 4
**
Received HDAC inhibitor [%]
Received chemo therapy [%]
Received ONC201 [%]
Received immuno therapy [%]
Proportion of Locations
Cluster
no
yes
Brainstem
Spine
Thalamus

**Sup. Figure 2: Uniform Manifold Approximation and Projection (UMAP) of cell types and identification and analysis of patient clusters.**

(A) All classified cell types were found in complete patient cohort (n = 76) including primary tumor site, metastatic and adjacent healthy tissue. Malignant (H3K27M+) cluster, defines DMG tumor cells and is clearly separated from other groups. Myeloid, lymphoid and immune other (including both myeloid and lymphoid), make the middle cluster, whereas the third cluster harbor healthy neurons, astrocytes and oligodendrocyte.

(B) Correlation analysis regarding a univariate association between the cell type contribution of malignant (top) and immune (bottom) cells with overall survival (OS) for each of the four patient clusters identified. We obtained the following spearman correlation coefficients (rho) (malignant; myeloid): **cluster 1**: NA, 2 patients, **cluster 2**: rho = 0, p-value = 1; rho = -0.4, p-value = 0.6; **cluster 3**: rho = -0.55 , p-value = 0.16 ; rho = 0.89 , p-value = 0.002; **cluster 4**: rho = 0.003, p-value = 0.99; rho = 0.06 , p-value = 0.77 ; Apart from cluster 3 (significantly ** larger OS with larger fraction of Myeloid cells), no significant ($p < 0.05$) associations were observed. (** $p < 0.01$)

(C) Characterization of the four patient clusters with respect to treatments received (HDAC inhibitors, chemo therapy, ONC201, or immunotherapy) as well as anatomical location. No specific associations between selected clusters and these factors were observed implying that clusters were mixed regarding patients receiving or not receiving the treatment. Note that cluster 1 comprised only 2 patients. Cluster 3 comprised exclusively brainstem DMGs.

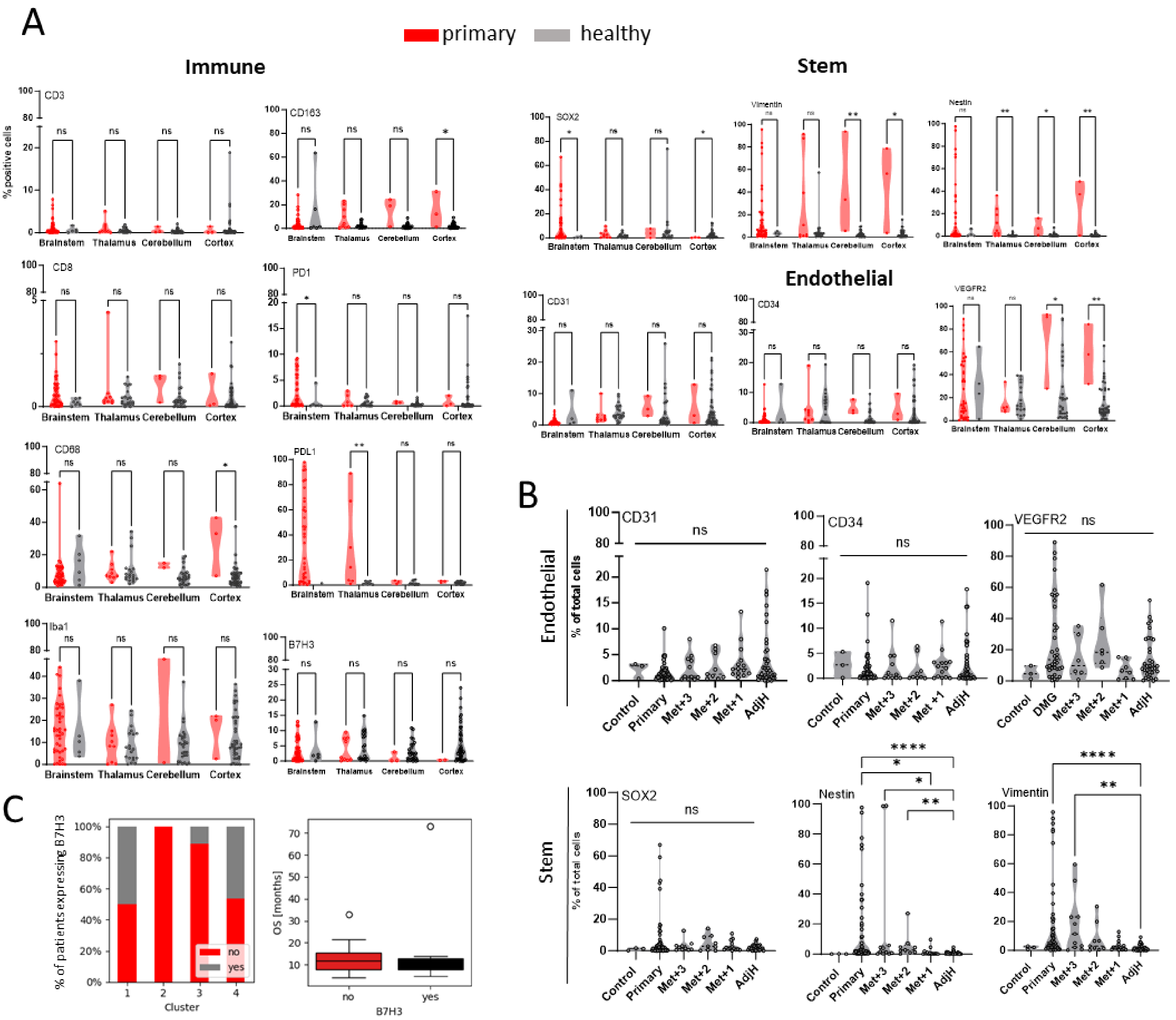
A
primary
healthy
Immune
Stem
Endothelial
% positive cells
CD3
CD163
CD8
PD1
CD68
PDL1
Iba1
B7H3
SOX2
Vimentin
Nestin
CD31
CD34
VEGFR2
Brainstem Thalamus Cerebellum Cortex
ns
B
Endothelial
Stem
% of total cells
Control
Primary
DMG
Met+3
Met+2
Met+1
AdjH
C
% of patients expressing B7H3
Cluster
no
yes
OS [months]
B7H3

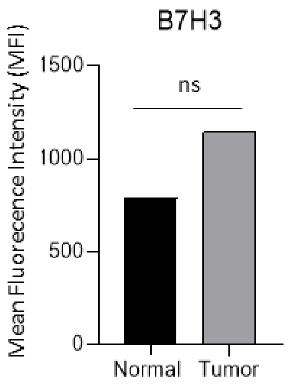
B7H3
ns
Mean Fluorecence Intensity (MFI)
1500
1000
500
0
Normal
Tumor

**Sup. Figure 3: Immune, stem and endothelial cell type mapping across sub anatomical brain locations and across different brain tumors and in DMG cohort primary, metastatic sites and adjacent and healthy brain tissue. B7H3 analysis amongst the cohort and FACS analysis of patient adjacent healthy (normal) and primary tumor tissue.**

(A) Comparison of cells positive for immune markers (CD3, CD8, CD68, Iba1, B7H3, PDL1, PD1), stem cell (SOX2, nestin, vimentin), and endothelial markers (CD31, CD34, VEGFR2) between primary brain tumors (red) and healthy brain tissues (gray) across different brain regions (brainstem, thalamus, cerebellum, cortex).

(B) Analysis of vascularization (CD31, CD34 and VEGFR2) and stem cell markers (SOX2, nestin and vimentin) across primary and metastatic sites compared to adjacent healthy and non-CNS tumor control patients.

(C) B7H3 expression was not associated with a specific cluster of patients as identified by hierarchical clustering based on cell type compositions (see Fig. 2D). FACS analysis of one DMG patient tumor and matched adjacent healthy sites for B7H3+ cells.

Mann-Whitney test for non-parametric pairwise comparison and Kruskal-Wallis test for non-parametric data for multiple group comparisons was used as statistical tests. (* $p < 0.05$, ** $p < 0.01$, *** $p < 0.001$, **** $p < 0.0001$)

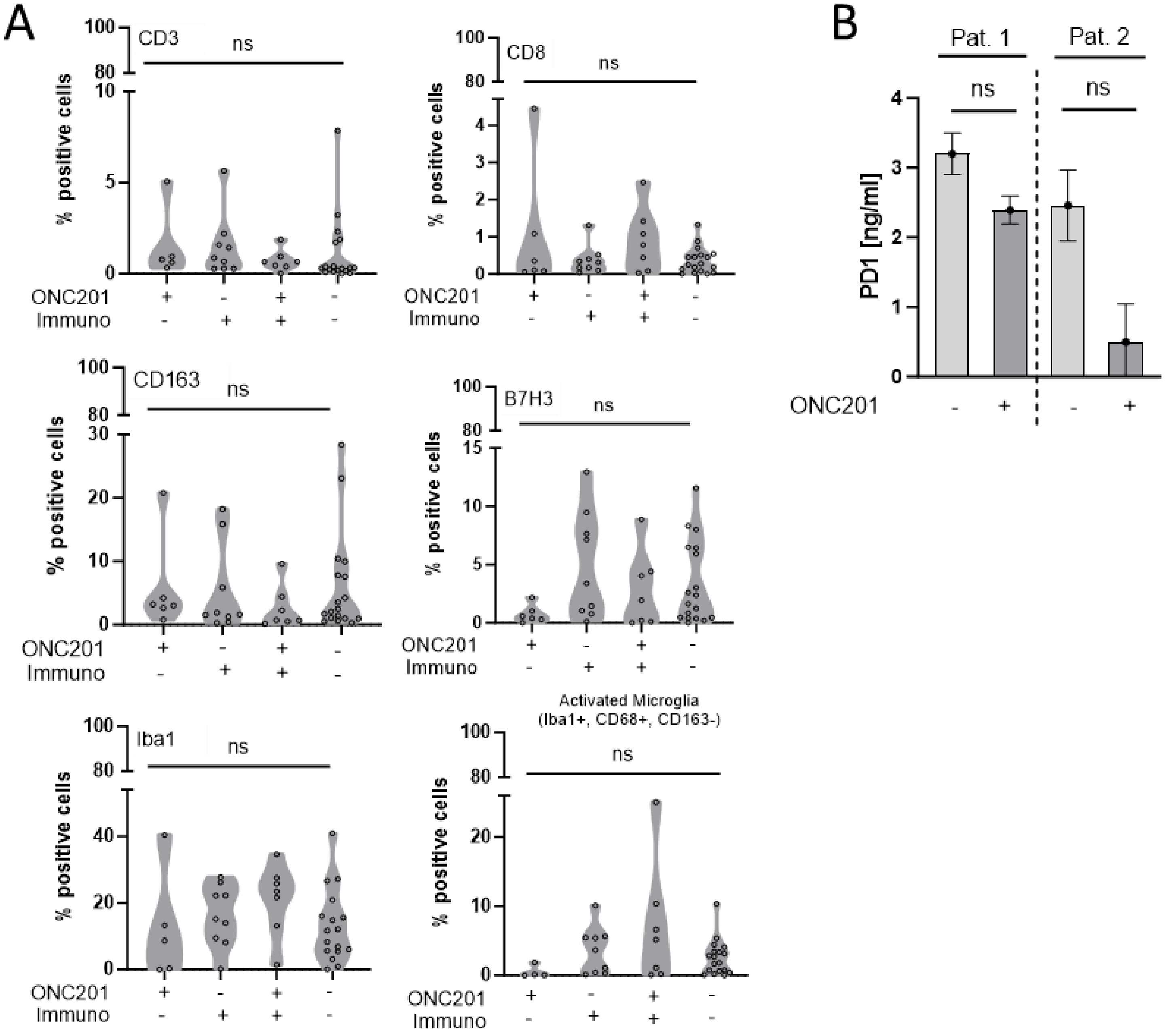
A
CD3
CD8
CD163
B7H3
Iba1
Activated Microglia (Iba1+, CD68+, CD163-)
ns
% positive cells
ONC201
Immuno
B
Pat. 1
Pat. 2
PD1 [ng/ml]
ONC201

**Sup. Figure 4: Immune marker mapping in DMG cohort primary sites and patient plasma in the context of treatment.**

(A) Comparison of percentage (%) cells positive for immune markers (CD3, CD8, CD163, B7H3, Iba1, activated Microglia) between patients treated with ONC201 (n = 5), Immunotherapy (n = 9), or combination (n = 7) compared to ONC201 and immune naïve (n = 17) patients.

(B) Plasma from patients receiving ONC201 were collected pre and post therapy and assessed additionally for immune suppression marker PD1.

Mann-Whitney test for non-parametric pairwise comparison and 2way ANOVA with Tukey's post-hoc test was applied for pairwise comparisons. (* $p < 0.05$, ** $p < 0.01$, *** $p < 0.001$, **** $p < 0.0001$)

| Treatment received | CD3 [%] | CD8 [%] | CD163 [%] | B7H3 [%] | Iba1 [%] | Activated microglia [%] |
|---|---|---|---|---|---|---|
| **ONC201** | 1.6 ± 2.0 | 1.0 ± 1.7 | 5.8 ± 7.4 | 0.7 ± 0.8 | 12.5 ± 16.6 | 0.5 ± 0.9 |
| **Immunotherapy** | 1.5 ± 1.7 | 0.4 ± 0.4 | 5.2 ± 6.9 | 4.9 ± 4.6 | 16.1 ± 9.2 | 3.6 ± 3.4 |
| **Combination** | 0.7 ± 0.6 | 0.9 ± 0.9 | 2.6 ± 3.4 | 2.8 ± 3.3 | 21.1 ± 10.8 | 6.9 ± 8.8 |
| **Other treatments** | 1.1 ± 1.9 | 0.4 ± 0.4 | 6.0 ± 8.0 | 3.4 ± 3.5 | 13.1 ± 10.8 | 2.6 ± 2.6 |

**Sup. Table 3**: **Percentage of immune marker positive cells per treatment group.**

Assessment of percentage of immune marker positive cells in ONC201 treated patients (n = 5), immunotherapy (n = 9), combination (n = 7) and treatment naïve (n = 17) resulted in non-significant increased immune cells for B7H3, Iba1 and activated microglia in combinational treated patients. Table showing mean and standard deviation (SD).

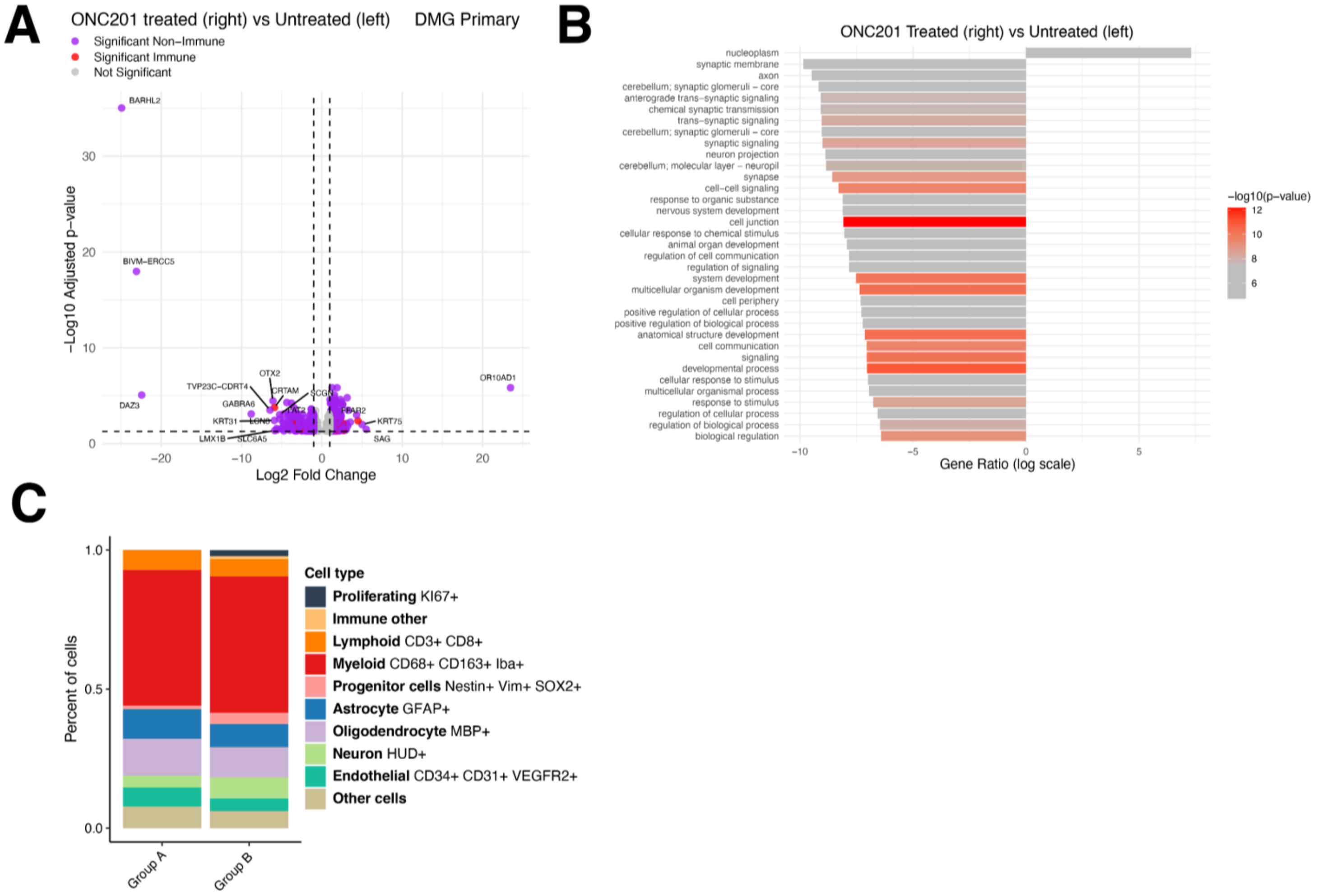
A
ONC201 treated (right) vs Untreated (left)
DMG Primary
Significant Non−Immune
Significant Immune
Not Significant
BARHL2
BIVM−ERCC5
DAZ3
OTX2
TVP23C−CDRT4
CRTAM
GABRA6
SCGN
KRT31
LMX1B
SLC6A5
OR10AD1
KRT75
SAG
−Log10 Adjusted p−value
Log2 Fold Change
B
ONC201 Treated (right) vs Untreated (left)
nucleoplasm
synaptic membrane
axon
cerebellum; synaptic glomeruli − core
anterograde trans−synaptic signaling
chemical synaptic transmission
trans−synaptic signaling
cerebellum; synaptic glomeruli − core
synaptic signaling
neuron projection
cerebellum; molecular layer − neuropil
synapse
cell−cell signaling
response to organic substance
nervous system development
cell junction
cellular response to chemical stimulus
animal organ development
regulation of cell communication
regulation of signaling
system development
multicellular organism development
cell periphery
positive regulation of cellular process
positive regulation of biological process
anatomical structure development
cell communication
signaling
developmental process
cellular response to stimulus
multicellular organismal process
response to stimulus
regulation of cellular process
regulation of biological process
biological regulation
Gene Ratio (log scale)
−log10(p−value)
C
Percent of cells
Cell type
Proliferating KI67+
Immune other
Lymphoid CD3+ CD8+
Myeloid CD68+ CD163+ Iba+
Progenitor cells Nestin+ Vim+ SOX2+
Astrocyte GFAP+
Oligodendrocyte MBP+
Neuron HUD+
Endothelial CD34+ CD31+ VEGFR2+
Other cells
Group A
Group B

**Sup. Figure 5: Transcriptomic in ONC201-treated versus untreated DMG cases and cellular differences across RNA groups A and B.**

(A) Volcano plot showing the differential gene expression analysis between ONC201-treated (right) and untreated (left) primary DMG samples. Significance (a<0.05) and fold-change in expression represented a log10 and log2 scale, respectively. Immune-related genes colored in red.

(B) Functional enrichment analysis (g:Profiler) from differentially expressed genes identified in (A) comparing ONC201-treated and untreated DMG samples. Synaptic and neuronal-related pathways, including “synaptic membrane,” “chemical synaptic transmission,” and “nervous system development,” were enriched in ONC201-untreated samples.

(C) Spatial proteomic cell type composition comparing RNA groups A and B. No significant differences in cell populations were observed between the two groups.
